# Structure-aware Graph Learning Predicts RNA Editability Across Tissues and Species

**DOI:** 10.64898/2026.02.02.703375

**Authors:** Zohar Rosenwasser, Michael Levitt, Erez Y. Levanon, Gal Oren

## Abstract

Programmable A-to-I RNA editing using endogenous ADAR enzymes is emerging as a therapeutic strategy, but editability remains difficult to predict because ADAR recognition depends on double-stranded RNA geometry and stability rather than sequence alone. We present ADAREDIT, a structure-explicit graph-attention framework that represents each dsRNA substrate as a nucleotide graph with backbone and base-pair edges. The framework includes a baseline model and a bio-aware model, with the latter augmenting this representation with typed interactions and a motif-sensitive sequence branch. We trained and evaluated both models on high-confidence inverted *Alu* duplexes (*n* = 884) with secondary structures predicted by RNAfold and editing levels measured across 8,603 GTEx RNA-seq samples spanning 47 tissues. Across five tissue contexts, the baseline and bio-aware models achieved strong held-out performance (test F1 = 0.814–0.869, AUROC = 0.869–0.933) and outperformed a matched structure-string baseline on the Liver split. The same graph representation retained predictive ability in evolutionarily distant non-*Alu* species (sea urchin, acorn worm, and octopus), suggesting conserved principles of ADAR substrate recognition. Finally, attention profiles and *in silico* mutagenesis recapitulated known biochemical constraints, including suppression by an upstream guanosine, and revealed longer-range asymmetric structural influences on editing. Because *Alu* duplexes are edited predominantly by ADAR1, ADAREDIT is geared primarily to the ADAR1 regime. ADAR1 is particularly relevant to therapeutic editing given its broad tissue expression.

The sources of this work are available at our repository: https://github.com/Scientific-Computing-Lab/AdarEdit

## 1 Introduction

Adenosine-to-inosine (A-to-I) RNA editing catalyzed by ADAR enzymes (Adenosine Deaminase Acting on RNA) represents a critical post-transcriptional mechanism that expands proteomic diversity and regulates immune responses Bass (2002); Nishikura (2010); Eisenberg & Levanon (2018). ADAR enzymes selectively deaminate adenosines within double-stranded RNA (dsRNA) structures, converting them to inosines that are interpreted as guanosines during translation. In mammals, ADAR1 and ADAR2 exhibit different tissue-specific expression patterns and substrate preferences, with ADAR1 predominantly editing repetitive *Alu* elements in untranslated regions Levanon et al. (2004); Bazak et al. (2014), while ADAR2 targets mainly specific coding sequences with functional consequences Tan et al. (2017); Nishikura (2016). Dysregulated editing has been implicated in cancer Han et al. (2015); Silvestris et al. (2019), neurological disorders Kawahara et al. (2004); Srivastava et al. (2017), and autoimmune conditions. Beyond its role in various biological pathways, ADAR presents a promising avenue for therapeutic RNA modification. Programmable ADAR-based editing platforms leverage synthetic guide RNAs to direct endogenous ADAR enzymes to specific target sites, enabling site-directed A-to-I conversion for correcting pathogenic mutations Qu et al. (2019); Merkle et al. (2019); Reautschnig et al. (2025). The success of such strategies, however, critically depends on understanding the sequence and structural features that govern ADAR substrate selection.

Despite extensive characterization of ADAR biochemistry, computational prediction of editing sites remains challenging due to the complex interplay between sequence context and RNA secondary structure (Appendix A). ADAR substrate recognition requires both local sequence motifs – including the well-established 5’ neighbor preference against guanosine Eggington et al. (2011); Lehmann & Bass (2000) – and global structural features such as stem length, loop geometry, and base-pairing stability. Early computational approaches relied on manually engineered features including structural parameters (loop length, stem stability, distance to junction) and sequence motifs (nucleotide composition, k-mer frequencies), achieving moderate accuracy Ouyang et al. (2018); Liu et al. (2021). The emergence of deep learning introduced convolutional neural networks that learned sequence patterns automatically, exemplified by EditPredict Wang et al. (2021), which processes linear sequence windows to classify editing sites. More recently, transformer architectures have been applied to RNA editing prediction: Helix Cao et al. (2025) employs a structure-aware attention mechanism that integrates base-pairing probability matrices directly into transformer layers, achieving a Spearman correlation of 0.84 for on-target guide-RNA editing efficiency, a distinct guide-optimization task that we discuss further in Appendix A; previous work evaluated RNA-FM (a masked language model trained on 23 million sequences Chen et al. (2022)) fine-tuned for editing prediction, and ADAR-GPT Rosenwasser et al. (2026) (continual fine-tuning of GPT-4o-mini on 201-nucleotide sequence windows), achieving competitive performance (F1=0.763 for ADAR-GPT) on liver tissue data. However, while Helix incorporates secondary structure through attention mechanisms, these approaches – including EditPredict, RNA-FM, Helix, and ADAR-GPT – ultimately operate on linear sequence representations rather than explicit graph-based encodings of dsRNA topology, limiting their ability to fully capture the relational nature of base-pairing that determines ADAR binding and catalytic efficiency.

Graph neural networks (GNNs) offer a principled framework for encoding RNA structure Wu et al. (2020); Zhang et al. (2021); Bongini et al. (2022) by representing molecules as graphs where nucleotides are nodes and relationships – both sequential backbone connectivity and Watson-Crick/wobble base pairing – are edges. This representation naturally captures the dsRNA topology essential for ADAR recognition while enabling models to learn multi-scale patterns spanning local nucleotide motifs to global stem-loop architecture. Here, we introduce ADAREDIT, a structure-explicit graph-attention framework Velickovic et al. (2017) comprising baseline and bio-aware models that integrate sequence and structure through explicit graph construction from RNA secondary structure predictions. Unlike sequence-only models, both ADAREDIT models encode base-pairing relationships directly as graph edges, allowing attention mechanisms to weight the importance of structural neighbors dynamically; the bio-aware model additionally incorporates explicit stem-loop geometry features. The bio-aware model additionally includes a parallel sequence branch – a convolutional neural network that captures local motifs known to influence ADAR substrate selection – and fuses the graph and sequence representations for final classification. Trained on high-confidence *Alu*-derived substrates across multiple human tissues and evolutionarily distant species, ADAREDIT outperforms a matched structure-string baseline, retains predictive ability across tissue contexts and non-mammalian species, and supports model interpretation through attention analysis and *in silico* mutagenesis that recapitulate known ADAR-associated patterns while highlighting additional positional and structural sensitivities in model predictions.

## 2 Results

### 2.1 Data construction

We constructed two complementary datasets to evaluate graph-based RNA editing prediction:

### Human *Alu*-derived substrates (Figure 1A)

Following the protocol established in previous work Rosenwasser et al. (2026), we systematically identified stable dsRNA substrates by scanning human untranslated regions (UTRs) for the closest pairs of oppositely oriented *Alu* elements, yielding 905 high-confidence *Alu* pairs. Restricting candidate adenosines to those with *>* 100-read coverage leaves 884 unique substrate structures, which constitute the benchmark used throughout. For each pair, we predicted secondary structure using RNAfold Lorenz et al. (2011) and extracted editing levels from GTEx RNA-seq data (8,603 samples across 47 tissues) Lonsdale et al. (2013). The *Alu*-pair duplex sequences were reconstructed from the hg38 reference genome. Because reference-based reconstruction could in principle be affected by donor polymorphisms, we quantified both common population variation across the duplexes and donor-level variants at the labeled adenosines (Supplementary Fig. S4). Editing-mimicking donor variants overlapped only 0.151% of assayed sites and were label-balanced, indicating that donor genomic variation is unlikely to bias the A-to-I labels. The reconstructed duplexes form extended, stably paired dsRNA structures. Across the 884 substrates, a median of 81.2% of nucleotides are base-paired in the RNAfold structure. Among the 865 pairs that met the genomic-context folding length criterion, 94.6% retained an inter-arm paired fraction *≥*0.5 when re-folded with *±*200 bp of flanking hg38 sequence (Supplementary Fig. S6a,b). This stability is consistent with ADAR’s preference for extended double-stranded RNA Polson & Bass (1994); Bass (2002); Nishikura (2010). Each example corresponds to a single adenosine position within a predicted dsRNA duplex. Adenosines were classified as edited (*≥*15% editing level, *>* 100 read coverage) or non-edited (*<*1% editing, *>* 100 reads), with intermediate sites (1–15%) excluded to maintain binary classification. These intermediate sites constitute 25.7–34.9% of adenosines across the five tissues; in the Supplementary (Fig. S1, Table S1) we show that the models assign these held-out intermediate sites graded scores in proportion to their editing level, and that discrimination is robust to relaxing the cutoff (positive thresholds of 5%, 10%, and 15%). We created five tissue-specific datasets – Brain Cerebellum, Artery Tibial, Liver, Muscle Skeletal, and Combined (integrating all tissues) – where the RNA sequence and predicted structure remain identical across tissues, but editing levels and resulting classifications vary according to the measured editing levels from GTEx in each tissue. These five tissues were chosen to span the extremes of the ADAR1/ADAR2 expression axis rather than by dataset size (Supplementary Fig. S2): Liver is strongly ADAR1-dominant and is also the most accessible tissue for RNA-therapeutic delivery, making an ADAR1-oriented model of particular translational interest; Artery Tibial has the highest ADAR2 (*ADARB1*) expression of any GTEx tissue; Brain Cerebellum expresses both isoforms highly and also expresses the brain-enriched ADAR3, which has been proposed to negatively regulate editing; this tissue also shows the highest *Alu* editing index across tissues Roth et al. (2019); and Muscle Skeletal has the lowest ADAR1 expression and the lowest *Alu* editing index Roth et al. (2019). The Combined dataset pools editing sites across all 47 GTEx tissues, providing broad transcriptome-wide coverage. Because the same *Alu* duplex recurs across tissues and across sites, we assign complete *Alu* pairs rather than individual adenosine positions to disjoint train/validation/test splits (64:16:20) using a single assignment shared across all tissues, so that no *Alu* pair appears in more than one split and, in particular, never in one tissue’s training set and another tissue’s test set. Class balancing by downsampling to equal numbers of edited and non-edited adenosines is applied to the dataset as a whole before splitting; because whole *Alu* pairs are assigned to a single partition, the positive rate varies between splits (45.6–56.5%). Dataset sizes are detailed in Table 1 (Appendix B).

**Figure 1:**
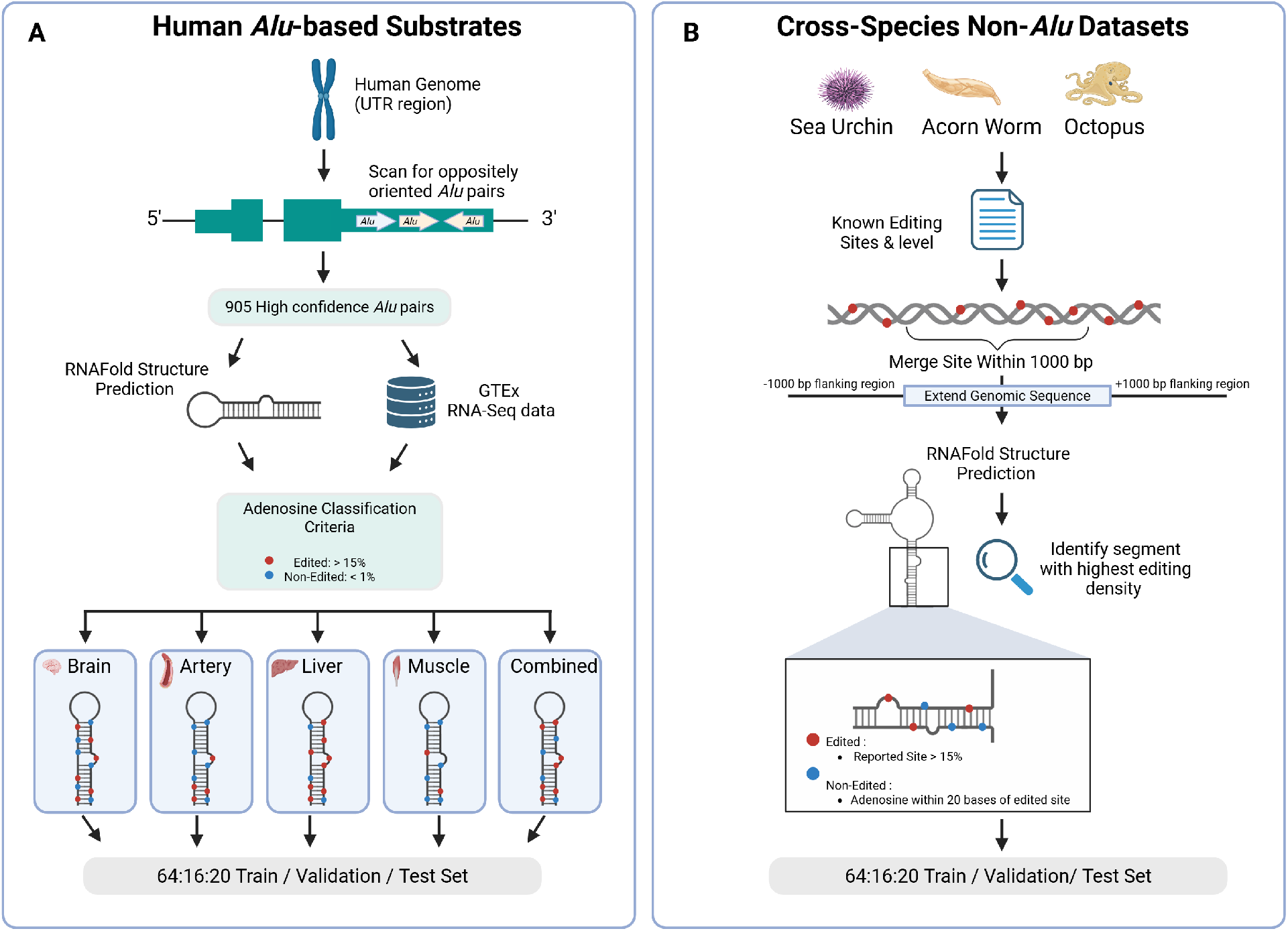
Schematic overview of data construction pipelines. (A) Construction of human *Alu* derived datasets. Candidate inverted *Alu* pairs were identified in UTRs (*n* = 905). After retaining candidate adenosines with *>* 100-read coverage, 884 distinct duplex structures were represented in the benchmark. For each pair, the secondary structure was predicted using RNAfold. Editing levels were obtained from GTEx RNA-seq data across varying tissues. Adenosines were classified as edited (*≥*15% level, *>* 100 coverage, red) or non-edited (*<*1% level, *>* 100 coverage, blue), creating five tissue-specific datasets where labels vary according to tissue-specific measurements while the sequence and structure remain constant. (B) Construction of cross-species non-*Alu* datasets (sea urchin, acorn worm, octopus). Known editing sites were clustered within 1,000 bp and extended with flanking regions. Following RNAfold structure prediction, the segment containing the largest number of reported editing sites was selected for each region. Candidate operational negatives were unreported adenosines collected within 20 bp of any source-cataloged editing site (blue), whereas benchmark positives were cataloged sites with editing levels *>* 15% (red). Class balancing was performed before complete *Alu* pairs (human) and editing clusters (species) were assigned to disjoint train/validation/test splits (64:16:20).

**Table 1:** Comprehensive performance metrics across all settings. Per-tissue models (each trained on one tissue and evaluated on all five held-out test tissues), cross-species transfer, and the joint multi-tissue model. For every setting we report F1, precision, recall, AUROC and AUPRC for the bio-aware (Bio) and baseline (Base) encoders, together with the decision threshold selected on validation; for the joint block, Bio and Base denote the joint bio-aware and joint baseline models. *N_train_*/*N_test_* denote dataset sizes, and bold marks the higher performance metric within each Bio/Base pair; values that are equal after rounding are left unbolded.

| Context | | Size ($N$) | | F1 Score | | Precision | | Recall | | AUROC | | AUPRC | | Threshold | |
| --- | --- | --- | --- | --- | --- | --- | --- | --- | --- | --- | --- | --- | --- | --- | --- |
| Train Set | Test Set | Train | Test | Bio | Base | Bio | Base | Bio | Base | Bio | Base | Bio | Base | Bio | Base |
| Artery Tibial | Artery Tibial | 15.3k | 4.8k | 0.858 | **0.867** | 0.831 | **0.834** | 0.888 | **0.902** | 0.918 | **0.924** | 0.923 | **0.926** | 0.400 | 0.450 |
|  | Brain Cer. | 15.3k | 4.6k | 0.864 | **0.871** | 0.815 | **0.824** | 0.919 | **0.925** | 0.924 | **0.928** | 0.927 | 0.927 | 0.400 | 0.450 |
|  | Liver | 15.3k | 4.2k | 0.856 | **0.864** | 0.822 | **0.827** | 0.893 | **0.905** | 0.910 | **0.913** | **0.929** | 0.925 | 0.400 | 0.450 |
|  | Muscle Skel. | 15.3k | 1.4k | 0.832 | **0.833** | **0.753** | 0.751 | 0.929 | **0.935** | **0.907** | 0.899 | **0.919** | 0.907 | 0.400 | 0.450 |
|  | Combined | 15.3k | 4.9k | 0.849 | **0.857** | **0.798** | 0.795 | 0.907 | **0.930** | 0.916 | **0.925** | 0.923 | **0.924** | 0.400 | 0.450 |
| Brain Cer. | Artery Tibial | 15.6k | 4.8k | 0.861 | **0.862** | **0.873** | 0.851 | 0.850 | **0.874** | **0.928** | 0.927 | **0.936** | 0.930 | 0.250 | 0.375 |
|  | Brain Cer. | 15.6k | 4.6k | 0.863 | **0.869** | **0.849** | 0.837 | 0.878 | **0.903** | 0.927 | **0.930** | **0.931** | 0.929 | 0.250 | 0.375 |
|  | Liver | 15.6k | 4.2k | 0.856 | **0.864** | **0.860** | 0.840 | 0.852 | **0.889** | 0.912 | **0.916** | **0.929** | 0.928 | 0.250 | 0.375 |
|  | Muscle Skel. | 15.6k | 1.4k | 0.842 | **0.845** | **0.792** | 0.776 | 0.899 | **0.928** | 0.904 | **0.905** | **0.918** | 0.911 | 0.250 | 0.375 |
|  | Combined | 15.6k | 4.9k | 0.857 | **0.863** | **0.830** | 0.819 | 0.885 | **0.912** | 0.927 | **0.931** | **0.935** | 0.934 | 0.250 | 0.375 |
| Liver | Artery Tibial | 12.5k | 4.8k | 0.830 | **0.832** | **0.845** | 0.835 | 0.815 | **0.829** | 0.902 | **0.904** | **0.914** | 0.909 | 0.375 | 0.525 |
|  | Brain Cer. | 12.5k | 4.6k | **0.843** | 0.837 | **0.840** | 0.822 | 0.846 | **0.853** | **0.913** | 0.907 | **0.921** | 0.909 | 0.375 | 0.525 |
|  | Liver | 12.5k | 4.2k | 0.854 | 0.854 | **0.854** | 0.837 | 0.855 | **0.872** | **0.910** | 0.909 | **0.927** | 0.924 | 0.375 | 0.525 |
|  | Muscle Skel. | 12.5k | 1.4k | **0.843** | 0.825 | **0.794** | 0.758 | 0.898 | **0.905** | **0.908** | 0.897 | **0.919** | 0.904 | 0.375 | 0.525 |
|  | Combined | 12.5k | 4.9k | 0.847 | **0.852** | **0.819** | 0.818 | 0.876 | **0.889** | 0.917 | **0.922** | 0.924 | **0.926** | 0.375 | 0.525 |
| Muscle Skel. | Artery Tibial | 4.7k | 4.8k | **0.798** | 0.788 | **0.837** | 0.833 | **0.762** | 0.747 | 0.871 | 0.871 | **0.891** | 0.878 | 0.100 | 0.350 |
|  | Brain Cer. | 4.7k | 4.6k | 0.802 | **0.811** | **0.823** | 0.821 | 0.782 | **0.802** | 0.881 | 0.881 | **0.886** | 0.884 | 0.100 | 0.350 |
|  | Liver | 4.7k | 4.2k | 0.809 | **0.821** | 0.832 | **0.844** | 0.788 | **0.798** | 0.871 | **0.882** | **0.897** | 0.894 | 0.100 | 0.350 |
|  | Muscle Skel. | 4.7k | 1.4k | **0.816** | 0.814 | **0.787** | 0.777 | 0.847 | **0.855** | **0.878** | 0.869 | **0.895** | 0.876 | 0.100 | 0.350 |
|  | Combined | 4.7k | 4.9k | **0.821** | 0.820 | **0.821** | 0.815 | 0.822 | **0.826** | 0.885 | **0.888** | **0.890** | 0.888 | 0.100 | 0.350 |
| Combined | Artery Tibial | 15.3k | 4.8k | 0.827 | **0.853** | 0.852 | **0.855** | 0.804 | **0.852** | 0.898 | **0.918** | 0.910 | **0.921** | 0.250 | 0.475 |
|  | Brain Cer. | 15.3k | 4.6k | 0.837 | **0.859** | 0.838 | **0.845** | 0.836 | **0.873** | 0.907 | **0.924** | 0.912 | **0.926** | 0.250 | 0.475 |
|  | Liver | 15.3k | 4.2k | 0.837 | **0.865** | 0.853 | **0.856** | 0.822 | **0.875** | 0.894 | **0.919** | 0.915 | **0.932** | 0.250 | 0.475 |
|  | Muscle Skel. | 15.3k | 1.4k | 0.835 | **0.855** | **0.795** | 0.793 | 0.878 | **0.927** | 0.902 | **0.909** | 0.910 | **0.914** | 0.250 | 0.475 |
|  | Combined | 15.3k | 4.9k | 0.836 | **0.867** | 0.826 | **0.833** | 0.846 | **0.904** | 0.903 | **0.933** | 0.910 | **0.936** | 0.250 | 0.475 |
| *Cross-Species Generalization* |  |  |  |  |  |  |  |  |  |  |  |  |  |  |  |
| Combined | *S. purp.* | 15.3k | 1.5k | 0.719 | **0.736** | 0.620 | **0.663** | **0.855** | 0.826 | 0.749 | **0.767** | **0.687** | 0.673 | 0.250 | 0.475 |
| Combined | *P. flava* | 15.3k | 1.6k | 0.717 | **0.772** | 0.665 | **0.744** | 0.777 | **0.802** | 0.702 | **0.809** | 0.668 | **0.799** | 0.250 | 0.475 |
| Combined | *O. bimac.* | 15.3k | 1.7k | **0.781** | 0.778 | 0.734 | **0.783** | **0.834** | 0.773 | 0.803 | **0.816** | 0.807 | **0.823** | 0.250 | 0.475 |
| *S. purp.* | *S. purp.* | 4.9k | 1.5k | 0.734 | **0.797** | 0.634 | **0.722** | 0.872 | **0.888** | 0.797 | **0.860** | 0.772 | **0.831** | 0.375 | 0.400 |
| *P. flava* | *P. flava* | 4.8k | 1.6k | 0.756 | **0.791** | 0.722 | **0.726** | 0.794 | **0.870** | 0.810 | **0.823** | 0.820 | **0.831** | 0.100 | 0.475 |
| *O. bimac.* | *O. bimac.* | 4.8k | 1.7k | **0.831** | 0.809 | 0.789 | **0.831** | **0.878** | 0.788 | **0.872** | 0.865 | 0.885 | 0.885 | 0.325 | 0.600 |
| *Joint multi-tissue model (one model, five-way head)* |  |  |  |  |  |  |  |  |  |  |  |  |  |  |  |
| Joint (all) | Artery Tibial | 37.0k | 4.8k | **0.859** | 0.853 | 0.832 | **0.841** | **0.889** | 0.866 | **0.924** | 0.915 | **0.934** | 0.919 | 0.175 | 0.425 |
| Joint (all) | Brain Cer. | 37.0k | 4.6k | **0.863** | 0.855 | 0.821 | **0.846** | **0.910** | 0.864 | **0.926** | 0.917 | **0.932** | 0.918 | 0.125 | 0.425 |
| Joint (all) | Liver | 37.0k | 4.2k | **0.862** | 0.860 | **0.858** | 0.846 | 0.866 | **0.875** | **0.916** | 0.911 | **0.935** | 0.924 | 0.175 | 0.400 |
| Joint (all) | Muscle Skel. | 37.0k | 1.4k | **0.836** | 0.829 | **0.825** | 0.767 | 0.848 | **0.901** | **0.898** | 0.892 | **0.913** | 0.896 | 0.100 | 0.325 |
| Joint (all) | Combined | 37.0k | 4.9k | 0.854 | **0.856** | 0.818 | **0.821** | 0.893 | **0.894** | **0.924** | 0.919 | **0.931** | 0.919 | 0.125 | 0.375 |

### Cross-species non-*Alu* datasets (Figure 1B)

To test generalization beyond human *Alu* elements, we constructed datasets for three evolutionarily distant species lacking *Alu* repeats: sea urchin (*Strongylocentrotus purpuratus*), acorn worm (*Ptychodera flava*), and octopus (*Octopus bimaculoides*). For each species, we obtained documented editing sites and their measured editing levels from Zhang et al. Zhang et al. (2023). We merged editing sites located within 1,000 bp into discrete clusters containing more than five sites, and then extracted an extended genomic sequence spanning each cluster plus 1,000 bp flanking regions on both sides. For each extended sequence, we predicted the minimum free-energy secondary structure using RNAfold Lorenz et al. (2011) and identified the segment containing the largest number of reported editing sites. Adenosines with reported editing levels *>* 15% and at least 100 supporting reads were labeled as positive. Unreported adenosines within 20 bp of any source-catalogued editing site were used as operational negatives. Because the sites used to collect candidate negatives were not required to exceed the 15% positive threshold, some operational negatives lie more than 20 bp from the nearest benchmark-positive site.

We tested sensitivity to this labeling rule by re-evaluating the fixed held-out predictions after excluding operational negatives at distances *≤* 5, *≤* 10, or *≤* 30 bp of a benchmark-positive site in the same dsRNA region and arm (Supplementary Fig. S5). Models were not retrained, and the validation-selected thresholds were unchanged. Each species dataset was class-balanced and partitioned into region-disjoint train, validation, and test sets (64:16:20).

### 2.2 ADAREDIT Model

#### A conceptual gap in modeling RNA editing

Many computational approaches to A-to-I RNA editing have operated on linear representations of RNA, either by modeling sequence windows directly or by augmenting them with engineered descriptors. This is not due to a lack of recognition that structural features matter – the importance of A:C mismatches, dsRNA stability, and stem-loop geometry has long been acknowledged – but rather because structural information is often incorporated as local annotations or summary features rather than as an explicit representation that preserves the relational topology of base pairing. However, ADAR-mediated editing is inherently a *structural* phenomenon: it depends on the formation, stability, and geometry of double-stranded RNA, rather than on sequence motifs alone.

As a result, prior models face an inherent limitation that is not merely architectural, but conceptual: learning from linearized representations can obscure the relational nature of base pairing, where nucleotides distant along the backbone become direct neighbors in the duplex and jointly shape ADAR access and activity. This motivates a structure-native representation that can directly encode both sequential and base-pairing relationships within a unified input.

We developed two graph neural network architectures to investigate the contribution of biology-aware feature engineering to RNA editing prediction. Both models represent RNA duplexes as graphs where nucleotides are nodes and edges encode both sequential connectivity and base-pairing relationships predicted from secondary structure (Figure 2). Each example corresponds to a single candidate adenosine, marked as the target node within the predicted duplex graph.

#### Baseline graph architecture

The baseline model employs a streamlined feature set (Figure 2, left panel): each node is encoded with an 8-dimensional vector comprising one-hot base identity (A, G, C, U/T, N), a binary pairing status flag, relative distance from the candidate editing site, and a target site indicator. Edges are constructed between sequential neighbors (i-to-i+1 backbone connectivity) and between base-paired nucleotides identified by parsing dot-bracket notation from RNAfold structure predictions Lorenz et al. (2011), creating a bidirectional graph that captures both linear sequence context and long-range structural constraints. Three stacked Graph Attention Network (GAT) layers with multi-head attention (4 heads per layer) process this graph, with each layer applying ReLU activation, batch normalization, and dropout (rate=0.2) for regularization. The attention mechanism dynamically learns which neighboring nucleotides – whether sequential or structurally paired – contribute most to editing prediction at each layer. After three rounds of message passing, global mean pooling aggregates node embeddings across the entire graph into a fixed-size representation, which is passed through a fully connected layer followed by sigmoid activation to produce the binary editing probability. This baseline architecture explicitly encodes RNA secondary-structure topology, distinguishing its input representation from sequence-window models. In the baseline, edges carry no feature vector (edge dimension = 0): backbone and base-pair edges are treated identically, so the model receives only graph topology with no edge-type signal. A full formal node/edge feature specification for each architecture is given in Supplementary Table S2.

**Figure 2:**
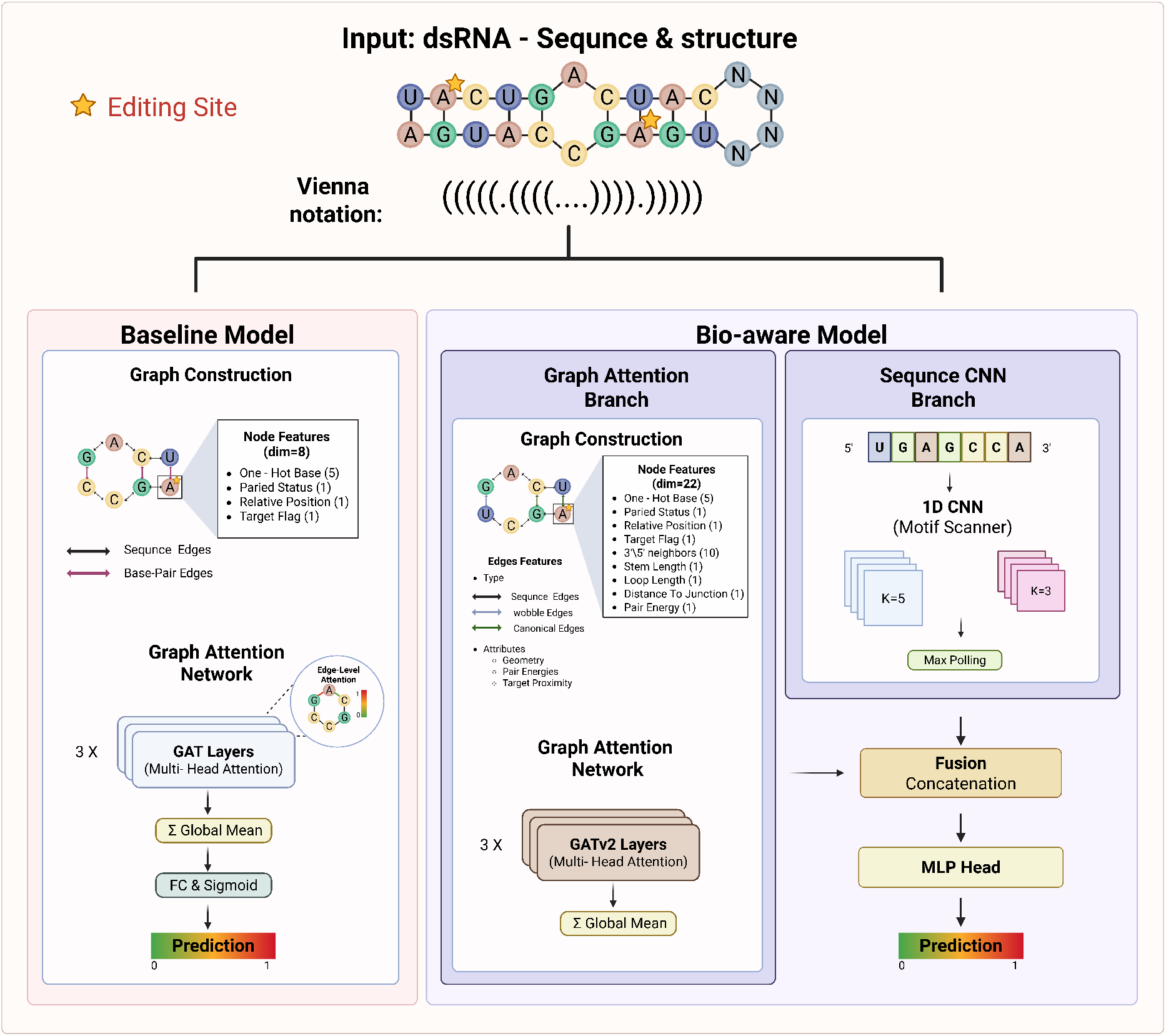
ADAREDIT architecture comparing baseline and bio-aware graph neural networks for RNA editing prediction. The figure illustrates the complete workflow for RNA editing site prediction using graph-based representations of double-stranded RNA structures. Top: Input representation showing dsRNA sequence and secondary structure with the candidate editing site (marked with star) and corresponding Vienna dot-bracket notation. Left (Baseline): Graph representation with 8-dimensional node features (base identity, pairing status, relative position, target flag) and edges connecting sequential neighbors (black) and base pairs (purple). Three GAT layers with multi-head attention, followed by global mean pooling and classification. Right (Bio-aware): Two-branch architecture: (1) Graph attention branch with enriched 22-dimensional node features (base identity, neighboring bases, structural geometry, and pair-energy scores) and typed backbone and base-pair edges; (2) Sequence CNN branch with 1D convolutions (K=3, K=5) capturing local motifs. Graph and sequence embeddings are fused through an MLP head for final prediction.

#### Bio-aware model enhancements

The bio-aware model is a deliberate biology-aware extension whose purpose is to test whether explicit biochemical and structural annotations add predictive value beyond the compact baseline graph, which already encodes dsRNA topology. The bio-aware model extends the baseline architecture by incorporating explicit biochemical knowledge at multiple levels (Figure 2, right panel). First, we distinguish sequential backbone edges from structural base-pair edges, including canonical Watson–Crick pairs (A–U, G–C) and wobble pairs (G–U). Each edge type receives a learned embedding (dimension = 6) that is concatenated with scalar edge attributes – including sequence/pair indicators, normalized sequence distance, stem length at both endpoints, loop length at both endpoints, distance to the nearest stem–loop junction, the pair-energy score defined below, and the relative distance of both endpoints from the target site – and passed through a small MLP to produce a 16-dimensional edge representation supplied to the GAT layers; these typed edge features are specified, alongside the baseline, in Supplementary Table S2. Second, we enrich node features beyond the baseline’s 8 dimensions by adding: one-hot encodings of the immediate 5’ and 3’ neighbors (10 dimensions total, capturing trinucleotide context known to influence ADAR selectivity), stem-loop geometry features (stem length, loop length, distance to junction), and a one-dimensional pair-energy score. Pair energy was represented by a nucleotide-identity score reflecting relative pairing strength (G–C, *−*3.4; A–U, *−*2.1; G–U, *−*1.0; and 0 for unpaired or other identities). Together, these features yield a 22-dimensional node representation combining local sequence, structural geometry, and pairing information. Third, we introduce a parallel sequence branch consisting of a 1D convolutional neural network that processes the tokenized RNA sequence independently of the graph: separate 3-mer and 5-mer convolutional filters (128 channels each, with padding to preserve length) capture local motifs, followed by masked max-pooling over sequence positions and projection to a 128-dimensional sequence embedding. Fourth, we optionally incorporate a global attention mechanism: a multi-head self-attention layer (4 heads) operates on the dense node embeddings after GAT processing, computing attention-weighted aggregation across all positions to capture long-range dependencies beyond the graph’s local message-passing horizon, yielding an additional global context vector. Finally, the graph-pooled embedding, sequence CNN embedding, and (if enabled) global attention embedding are concatenated and passed through a two-layer MLP classification head with ReLU activation and dropout. All structural information derives from RNAfold predictions of the secondary structure. The bio-aware model thus integrates graph topology, edge biochemistry, local sequence motifs, structural geometry, and optional global attention into a unified architecture, enabling it to learn multi-scale editing determinants while maintaining full differentiability for end-to-end training. This optional layer was disabled in all models reported here.

A critical aspect of both architectures is the position-centric graph encoding scheme. While multiple adenosines within the same duplex share identical RNA sequence and secondary structure, each candidate site generates a distinct graph representation. Node features encoding relative distance from the candidate editing site and the binary target indicator ensure that the feature values differ for each evaluated position, so each candidate adenosine yields a distinct position-centric graph representation even when several sites share the same underlying duplex topology. Leakage across splits, however, is prevented by the data split itself: complete *Alu* pairs are held out together (Data construction), rather than by these per-site representation differences.

### A single joint model across tissues

In addition to the per-tissue models, we trained joint versions of both encoders across all five tissue contexts. Each joint model uses the same encoder architecture as its per-tissue counterpart, but replaces the single binary output with a five-output multi-label head that predicts editing separately for Artery Tibial, Brain Cerebellum, Combined, Liver, and Muscle Skeletal. The model is trained with a masked binary cross-entropy loss, so that each site contributes only to the tissues in which it was measured. For the joint models, thresholds were optimized separately for each tissue output, and checkpoint selection was based on macro validation F1 across the observed outputs. This represents each candidate adenosine once while retaining tissue-specific outputs, and we compare the joint models against the separate per-tissue models in the Results (Figure 3D).

**Figure 3:**
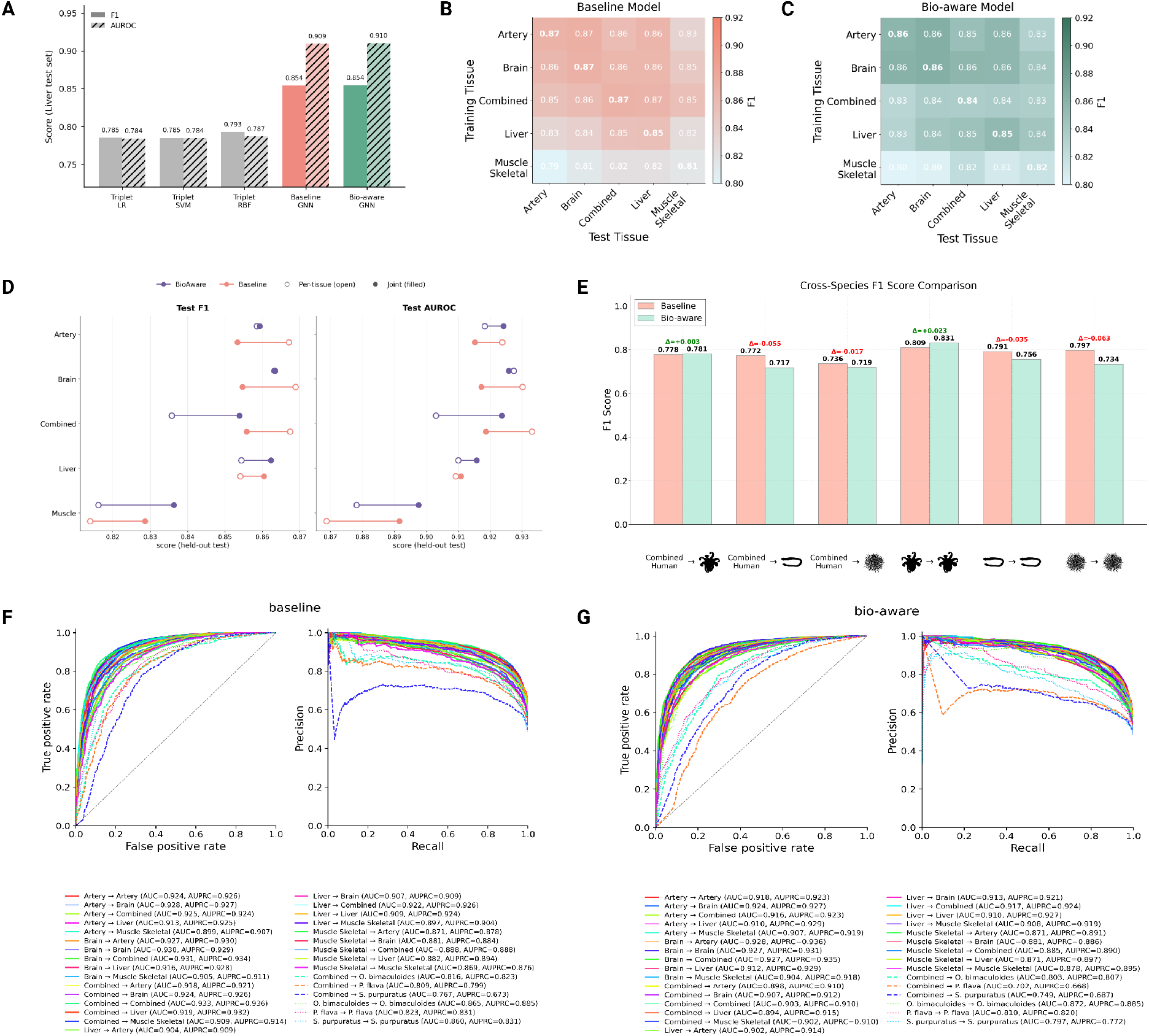
**Performance and generalization of graph-based RNA editing prediction models.**(A) F1 and AUROC on the held-out Liver test set for the target-centered structure-string Triplet baselines (logistic regression, linear SVM, and RBF-kernel SVM) and the two graph-based architectures (Baseline GNN, Bio-aware GNN), all evaluated on the identical split. (B,C) Cross-tissue F1 score heatmaps for the baseline (B) and bio-aware (C) ADAREDIT models. Rows indicate training tissue and columns indicate test tissue. Diagonal entries correspond to within-tissue evaluation; off-diagonal entries quantify cross-tissue generalization across distinct ADAR expression contexts. (D) Test F1 (left) and AUROC (right) of the joint multilabel model versus the five separate per-tissue models, shown for both the baseline (orange) and bio-aware (purple) encoders; for each tissue, open circles mark the per-tissue model and filled circles the joint model. (E) Cross-species F1 score comparison for (i) within-species training and evaluation (species*→*species) and (ii) transfer from the human Combined model to non-*Alu* species (Combined*→*species), shown for Strongylocentrotus purpuratus, Ptychodera flava, and Octopus bimaculoides. Bars denote baseline (pink) and bio-aware (green) performance; ΔF1 values are annotated. (F,G) ROC (left) and PR (right) curves for all cross-tissue and cross-species train–test combinations for the baseline (F) and bio-aware (G) models, illustrating model discrimination across diverse training and evaluation settings.

#### Model selection via F1-optimized threshold search

To select the best-performing model during training while accounting for the threshold-dependent nature of classification metrics, we employed an F1-optimized threshold search strategy. After each training epoch, we evaluated the model on the held-out validation set and computed performance metrics across a dense grid of 33 decision thresholds, linearly spaced from 0.1 to 0.9. For each threshold candidate, we calculated five classification metrics: accuracy, F1-score, precision (positive predictive value), recall (sensitivity), and specificity (true negative rate). Among all threshold-epoch combinations evaluated during training, we selected the model checkpoint that achieved the maximum F1-score, as this metric balances precision and recall – a critical consideration for RNA editing prediction where both false positives (mispredicted editing sites) and false negatives (missed genuine sites) carry biological and therapeutic consequences. This threshold-adaptive selection procedure was applied independently to each train*→*validation experiment, identifying the optimal operating point for each tissue or species context. The checkpoint and decision threshold are selected on the validation split only; the selected model is then evaluated once on the held-out, pair-disjoint test split, which is not used for checkpoint or decision-threshold selection, and all reported performance numbers are test-set metrics. All reported results are single training runs at a fixed seed (42). For threshold-independent evaluation, we additionally computed AUROC (area under the receiver operating characteristic curve) and AUPRC (area under the precision-recall curve), which assess discrimination performance across the entire spectrum of decision boundaries and provide complementary evidence of model quality independent of any specific operating point. In addition, analysis of the training logs for the per-tissue and within-species models and of the distribution of validation-selected decision thresholds (Appendix C) shows smooth loss convergence, stable validation-F1 trajectories, and validation-selected thresholds below 0.5 for most settings, indicating that the reported checkpoints are not isolated single-epoch outliers (Figure 7).

### 2.3 Performance comparison against existing methods

We first asked whether the explicit graph representation is warranted, or whether a much simpler model of the same fixed structure would suffice. As a structure-string baseline, we implemented a Triplet-SVM-style classifier, in the spirit of the local structure–sequence features introduced for microRNA-precursor classification Xue et al. (2005): each candidate adenosine is encoded by the identity of its surrounding bases together with the paired/unpaired pattern of their local three-nucleotide structural context within a *±*25-nt window, and classified with logistic-regression, linear-SVM, and RBF-kernel SVM heads. Such a model draws on the same RNAfold structure as ADAREDIT but compresses it into fixed local features, whereas the graph representation keeps base-pairing as explicit edges over the full duplex, enabling the network to learn from the spatial geometry of stem-loop architectures rather than from a linear string. On the held-out pair-disjoint Liver test split, the Triplet baselines reached F1 0.785 and AUROC 0.784 with logistic-regression and linear-SVM heads, and an RBF-kernel SVM on the same features performed similarly (F1 0.793, AUROC 0.787). The graph models performed better on the same split: the Baseline GNN reached F1 0.854 and AUROC 0.909, and the bio-aware model reached F1 0.854 and AUROC 0.910, corresponding to improvements of approximately +0.06–0.07 in F1 and +0.12–0.13 in AUROC relative to the Triplet baselines (Figure 3A). Earlier sequence-only predictors, the convolutional EditPredict, the fine-tuned foundation model RNA-FM, and the GPT-based ADAR-GPT report lower Liver F1 values (0.72, 0.71, and 0.763, respectively) Wang et al. (2021); Chen et al. (2022); Rosenwasser et al. (2026), but were evaluated on a separate, non-pair-disjoint sequence-window benchmark (fixed 201-nucleotide windows); we therefore cite them as published reference points rather than as a matched comparison (Appendix A).

### 2.4 Cross-tissue generalization reveals tissue-specific learning

To systematically assess within-context performance and cross-tissue behavior, we trained five independent baseline models, one for each human context (Liver, Brain Cerebellum, Artery Tibial, Muscle Skeletal, and Combined), and evaluated each model on all five test contexts, yielding a 5 *×* 5 matrix of 25 train*→*test evaluations (Figure 3B). This design revealed several patterns. First, within-tissue performance (diagonal elements of the matrix) achieved high F1 scores, with Liver*→*Liver at F1=0.854, Brain Cerebellum*→*Brain Cerebellum at F1= 0.869, and Artery Tibial*→*Artery Tibial at F1=0.867, indicating that models capture context-associated editing patterns when trained and tested on matched ADAR isoform contexts. Second, performance degraded when training and test sets had divergent ADAR expression profiles: for example, models trained on Liver (ADAR1-dominant) showed reduced performance when tested on Artery Tibial (ADAR2-dominant), with F1 dropping from 0.854 (Liver*→*Liver) to 0.832 (Liver*→*Artery Tibial), consistent with context-specific differences in editing patterns that may reflect differences in ADAR isoform expression. Third, Muscle Skeletal generally showed the lowest performance both as a training source and test target, likely reflecting the limited training data due to low ADAR1 expression and sparse editing activity in this tissue Roth et al. (2019). These cross-tissue experiments demonstrate that the baseline graph architecture successfully learns biologically meaningful patterns. ROC and precision–recall curves showed consistent discrimination across train*→*test tissue combinations (Figure 3F). To assess whether the GTEx-derived labels reflect reproducible editing measurements rather than a GTEx-specific processing artifact, we re-quantified *Alu* editing in an independent human cohort, RNAAtlas (216 samples, distinct sequencing centre and processing pipeline). Across 16,028 benchmark-selected sites (*<* 1% or *≥* 15% in GTEx) that were also measured in RNAAtlas, editing levels were strongly concordant with GTEx (Pearson *r* = 0.97, Spearman *ρ* = 0.93), with only one site changing edited/non-edited class between cohorts (Figure S3). We also tested whether performance depended primarily on substrate stability, given that the model receives a single RNAfold structure for each duplex. When test sites were stratified by substrate paired fraction, both architectures retained discrimination in the least-stable quartile (AUROC *≈*0.88–0.90 in most tissues; Muscle Skeletal, 0.83–0.84) and improved modestly in the most-stable quartile (Supplementary Fig. S6c,d).

We next asked whether explicit biochemical annotations improve prediction beyond the compact baseline graph. The bio-aware model augments the baseline topology with typed backbone and base-pair interactions, neighbouring-base context, stem–loop geometry, pair-energy scores, and a parallel sequence-CNN branch (Figure 2). Despite these additions, the two architectures performed similarly. Across the four tissue-specific held-out test sets, their F1 scores differed by no more than 0.009; in the Combined context, the baseline exceeded the bio-aware model by 0.032. Off-diagonal cross-tissue differences were also modest and generally favoured the baseline, whereas differences in AUROC and AUPRC were mixed (Figure 3B,C,F,G). This convergence across architectures indicates that the explicit graph topology provides the principal representational advantage: the compact baseline already makes relevant sequence and structural relationships accessible through message passing over backbone and base-pair edges, leaving limited incremental benefit from additional biochemical annotations in the per-context setting.

Component ablations on the Liver held-out test set further resolved the contributions of the bio-aware features (Figure S7). Removing neighbouring-base context produced the largest reduction in AUROC (ΔAUROC = *−*0.054), followed by scrambling the base-pairing partners (*−*0.024) and removing base-pair edges (*−*0.019). Removing the sequence-CNN branch or collapsing edge types had smaller effects (*−*0.005 and *−*0.002, respectively), whereas removing stem–loop geometry increased AUROC slightly in this run (+0.012). These results identify local sequence context and correct base-pair topology as the strongest contributors among the biology-inspired features. Notably, both sources of information are already accessible to the baseline through nucleotide identity and message passing over sequential and structural edges, explaining its strong performance despite its compact design. Because each ablation was trained once with a fixed seed, the smaller differences should be interpreted descriptively. Together, these findings establish the structure-native graph representation as the core strength of ADAREDIT: the baseline provides a parsimonious and high-performing implementation, while the bio-aware model offers a complementary framework for testing the incremental value of explicit biochemical annotations.

#### Joint training consolidates context-specific prediction in a single model

The analyses above used a separate model for each human context. We therefore asked whether shared multi-label training could replace this five-model setup while retaining context-specific outputs and exploiting signals shared across tissues. Because the *Alu*-duplex sequence and predicted structure of a site are identical across contexts whereas its editing status can vary, each candidate adenosine was represented once and processed by a shared encoder connected to a five-output multi-label head for Artery Tibial, Brain Cerebellum, Combined, Liver, and Muscle Skeletal (Figure 3D). A masked binary cross-entropy loss restricted each site’s contribution to contexts in which it was measured, without imputing unobserved labels. We trained joint versions of both encoders and compared them with their corresponding separate per-context models.

Joint training preserved overall predictive performance while replacing five separate encoders with a single shared model. The joint baseline achieved macro test F1 of 0.851 and AUROC of 0.911, compared with 0.854 and 0.913, respectively, for the separate baseline models. The joint bio-aware model improved over the corresponding separate bio-aware models, increasing macro F1 from 0.846 to 0.855, AUROC from 0.907 to 0.917, and AUPRC from 0.917 to 0.929. Performance was higher in four of the five contexts and was nearly unchanged in Brain Cerebellum. The largest improvement occurred in Muscle Skeletal, the smallest dataset, where F1 increased from 0.816 to 0.836 and AUROC from 0.878 to 0.898. These results are consistent with shared training reinforcing predictive sequence–structure signals across tissues, particularly in contexts with fewer observations.

Because the aggregate Combined label is derived from measurements across GTEx tissues and may provide partially redundant supervision, we also trained four-output joint models without Combined. Performance was retained or modestly improved: across the same four tissue outputs, macro F1 and AUROC changed from 0.849 and 0.909 to 0.858 and 0.914 for the baseline encoder, and from 0.855 and 0.916 to 0.856 and 0.920 for the bio-aware encoder, respectively (Supplementary Fig. S8). Thus, the performance of joint training was not contingent on the aggregate Combined output. Overall, a shared encoder can consolidate tissue-specific prediction into a single model while maintaining, and in some settings modestly improving, discrimination relative to separate per-context models.

### 2.5 Cross-species generalization suggests partially conserved structural principles

To assess whether ADAREDIT’s graph-based structural encoding captures editing determinants beyond human *Alu* elements, we evaluated models on three evolutionarily distant species lacking *Alu*: sea urchin (S. purpuratus), acorn worm (P. flava), and octopus (O. bimaculoides). These taxa span hundreds of millions of years of divergence and represent independent evolutionary trajectories of A-to-I editing, with editing occurring predominantly in species-specific repetitive elements rather than primate-derived SINEs. When trained and tested within each non-*Alu* species, both architectures achieved strong performance: baseline test F1 of 0.81 (octopus), 0.8 (sea urchin), and 0.79 (acorn worm), with corresponding bio-aware scores of 0.83, 0.73, and 0.76 (Figure 3E). These within-species results demonstrate that graph-based RNA structure encoding successfully captures ADAR substrate recognition patterns even in non-*Alu* contexts, supporting the conclusion that the learned predictive features are not restricted to human *Alu* substrates. More strikingly, models trained exclusively on human data (Combined dataset) transferred effectively to non-mammalian species: Combined*→*octopus reached F1=0.78, Combined*→*sea urchin F1=0.74, and Combined*→*acorn worm F1=0.77 for the baseline model; the corresponding bio-aware scores were 0.78, 0.72, and 0.72 (Figure 3E–G). Excluding operational negatives within 5, 10, or 30 bp of a benchmark-positive site from the fixed within-species held-out predictions increased F1, while AUROC remained stable or increased modestly, indicating that near-positive negatives do not inflate the reported species-level discrimination (Supplementary Fig. S5). While cross-species performance showed moderate degradation (F1 decreases of approximately 0.01–0.06 relative to within-species training), these results show substantial predictive performance and demonstrate that structural features learned from human *Alu* duplexes partially generalize to evolutionarily distant organisms. The performance reduction likely reflects genuine biochemical differences in ADAR substrate preferences evolved independently across lineages, while the substantial remaining predictive power indicates partial conservation of structural recognition features – dsRNA stability, structural constraints, and local sequence context – across deep evolutionary time. Comparing the two architectures across all six cross-species settings (Table 1), the baseline and bio-aware models perform comparably, with the baseline higher in four of six settings; both demonstrate effective cross-species transfer.

### 2.6 Transfer to selected protein-coding editing sites

To assess whether models trained exclusively on *Alu* elements generalize to functionally important editing sites in protein-coding transcripts, we evaluated performance on seven well-characterized coding-region targets across diverse genes: *AJUBA*, *BLCAP*, *TTYH2*, *NEIL1* (primarily ADAR1 substrates), and *FLNA*, *GRIA2*, *GRIA3* (primarily ADAR2 substrates), representing both constitutive and tissue-specific editing events (Figure 4). These targets were absent from training and provide a targeted out-of-domain scope test rather than a comprehensive coding-region benchmark. For each gene, we computationally identified the dsRNA structure encompassing the edited region by extracting transcript sequences flanking the target site and predicting secondary structure using RNAfold. We then catalogued all adenosines within these predicted dsRNA regions and classified them into two categories based on GTEx experimental editing data from three tissue contexts (Brain Cerebellum, Liver, Combined) as edited sites (*≥*15% editing) or non-edited sites (*<*1% editing), excluding intermediate sites (1–15% editing) as in the main analysis, creating evaluation sets for each gene-tissue combination. We then applied the separately trained Baseline and Bio-aware models and the two joint multilabel models introduced above, each trained on *Alu* pairs alone and run without retraining or fine-tuning, to score every adenosine, providing a targeted out-of-domain evaluation of all four model configurations (Figure 4A).

**Figure 4:**
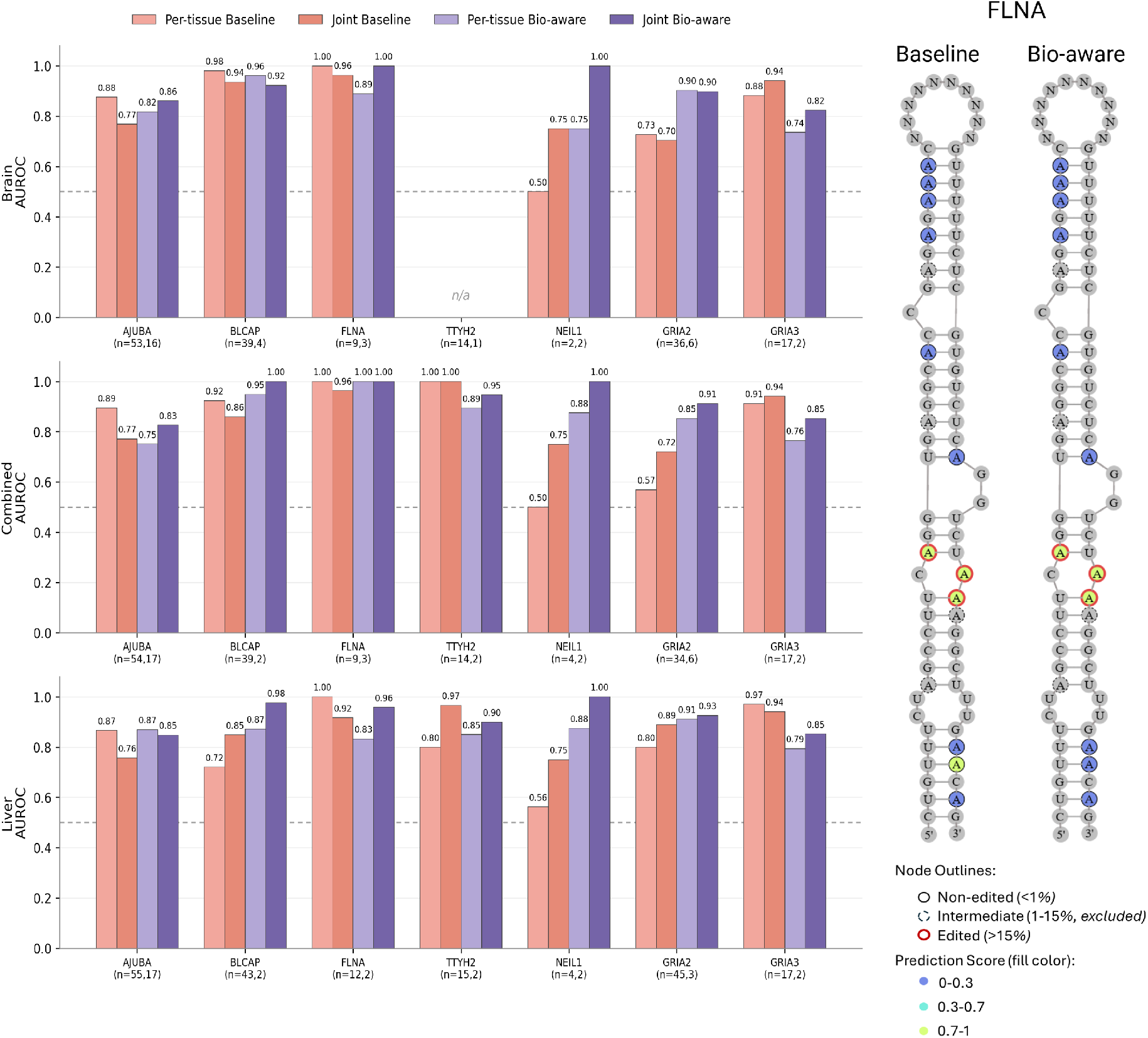
Transfer from *Alu* elements to protein-coding editing targets. (A) AUROC of the separately trained Baseline and Bio-aware models and their jointly trained counterparts across seven coding-region targets in Brain Cerebellum, Combined, and Liver. All models were trained exclusively on *Alu* pairs and applied without retraining or fine-tuning. Sites with *≥*15% editing were labelled edited, sites with *<*1% editing were labelled non-edited, and intermediate sites were excluded. Sample sizes are shown as (*n*_not_-_edited_, *n*_edited_); the dashed line denotes AUROC = 0.5, and n/a denotes fewer than two sites in either class. (B) Predicted editing scores across an example *FLNA* dsRNA structure for the Combined outputs of the Joint Baseline (left) and Joint Bio-aware (right) models. Node outlines indicate experimental annotations: solid, non-edited (*<*1%); dashed, intermediate (1– 15%, excluded); and red, edited (*≥*15%). Node fill indicates prediction score: dark blue, 0–0.3; cyan, 0.3–0.7; and green, 0.7–1.0.

We summarised performance using the area under the ROC curve (AUROC), which measures how reliably edited adenosines are ranked above non-edited adenosines independently of any decision threshold and is therefore directly comparable across the four models despite their different calibration. Because several targets contribute only a few sites per class, we report AUROC only where both classes contain at least two sites (*TTYH2* in Brain is therefore omitted) and interpret the per-gene values as indicative trends rather than precise estimates. The sparsest targets–notably *TTYH2* and *NEIL1*, with as few as 1–2 edited sites in some tissue contexts–are illustrative rather than statistically conclusive, and we draw no significance claims from them; a definitive per-gene assessment would require larger held-out coding sets.

All four models retained ranking ability on this coding-target scope test: across the seven targets and three tissue contexts, most AUROC values lay above the 0.5 chance level (Figure 4A). Because the training substrates are *Alu*-derived and thus predominantly ADAR1 targets, we expected the strongest transfer on ADAR1-associated coding sites. Consistent with this, the ADAR1-associated targets *AJUBA* and *BLCAP* reached AUROC ranges of 0.75–0.89 and 0.72–1.00, respectively, across contexts and models. Transfer to ADAR2-associated recoding sites was more variable but not absent: *FLNA* reached AUROC 0.83–1.00 in every context across the four models, although with only 2–3 edited sites per context, and *GRIA2*, the canonical Q/R target, was well separated by the Joint Bio-aware model (AUROC 0.90–0.93). Conversely, *GRIA3* was ranked more strongly by the baseline models, reaching AUROC 0.88–0.97 for the separately trained Baseline and 0.94 for the Joint Baseline model. Together, these results provide evidence that *Alu*-trained graph models can use transferable sequence–structure signal in selected coding substrates from both ADAR regimes, while also showing that this transfer remains target-and model-dependent.

An illustrative example from the *FLNA* target (Figure 4B) shows the full dsRNA structure with predictions from the Combined outputs of the Joint Baseline and Joint Bio-aware models overlaid on experimental editing annotations, demonstrating how both models correctly identify the majority of edited sites while maintaining specificity.

Overall, all four models ranked coding-site editing above chance across most targets, supporting the interpretation that graph-based structural encoding captures signals that are not solely *Alu*-specific. Performance varied across ADAR2-associated targets and model architectures, as may be expected from the predominantly ADAR1-like composition of the *Alu*-derived training set; nevertheless, the strong ranking of *FLNA* across all four models, of *GRIA2* by the Joint Bio-aware model, and of *GRIA3* by the baseline models indicates that transfer is not restricted to ADAR1-associated targets. Notably, these coding targets form markedly shorter–and presumably less stable–dsRNA duplexes than the long *Alu* inverted repeats: *FLNA*, for instance, is edited within a duplex spanning only *∼*85 nt, compared with a mean of 586 nt for the *Alu* substrates, yet the *Alu*-trained models ranked its edited adenosines strongly in every tissue context. These results suggest that transfer can extend to shorter coding-region dsRNA contexts, while broad recoding-site prediction remains outside the model’s primary validated regime.

### 2.7 Model interpretability

#### Interpretability through attention analysis of the baseline model

A central motivation for employing graph attention networks in ADAREDIT was to combine high predictive accuracy with biological interpretability. Graph attention mechanisms offer a framework for interpretability by revealing which neighboring nodes receive high weights during prediction Knyazev et al. (2019); Brody et al. (2021). While these patterns reflect learned associations rather than proven causal relationships, they can guide hypothesis generation when patterns align with established biochemistry. To understand what structural and sequence patterns the model learns independently – without explicit biological guidance – we performed all interpretability analyses on the baseline architecture rather than the bio-aware variant. The baseline model receives only graph topology (sequence edges and base-pairing edges) and minimal node features (base identity, pairing status, position), forcing it to discover editing determinants entirely from data through the attention mechanism. By analyzing baseline attention weights, we can identify which positional and structural features the model autonomously prioritizes, offering unbiased insights into learned editing determinants.

We conducted two complementary interpretability approaches (Figure 5A). First, to validate that attention weights contain genuine predictive signal rather than spurious patterns, we extracted edge-level attention coefficients from the last GAT layer and aggregated them into positional features representing the maximum attention at each position in a *±*50 nucleotide window centered on the candidate editing site. These attention-derived positional features were then used to train an independent XGBoost classifier Chen & Guestrin (2016) on the validation split and evaluate it on the duplex-disjoint test split, providing a supervised assessment of whether attention alone captures editing determinants Belinkov (2022). To identify which specific positions contribute most to predictions, we applied SHAP (SHapley Additive exPlanations) analysis Lundberg & Lee (2017) to the XGBoost model using the validation data, ranking features by their impact on model output. We then retrained the XGBoost classifier on the validation split using only the top 20 SHAP-identified features and repeated SHAP analysis to confirm that this reduced feature set retained interpretability. Second, to directly examine attention patterns without supervised intermediaries, we performed descriptive statistical analyses of mean attention across the *±*50 nucleotide window across all held-out test sites, stratifying by editing status (edited vs non-edited sites), nucleotide identity (A, U, G, C at each position), and structural context (base-paired vs unpaired/loop positions).

**Figure 5:**
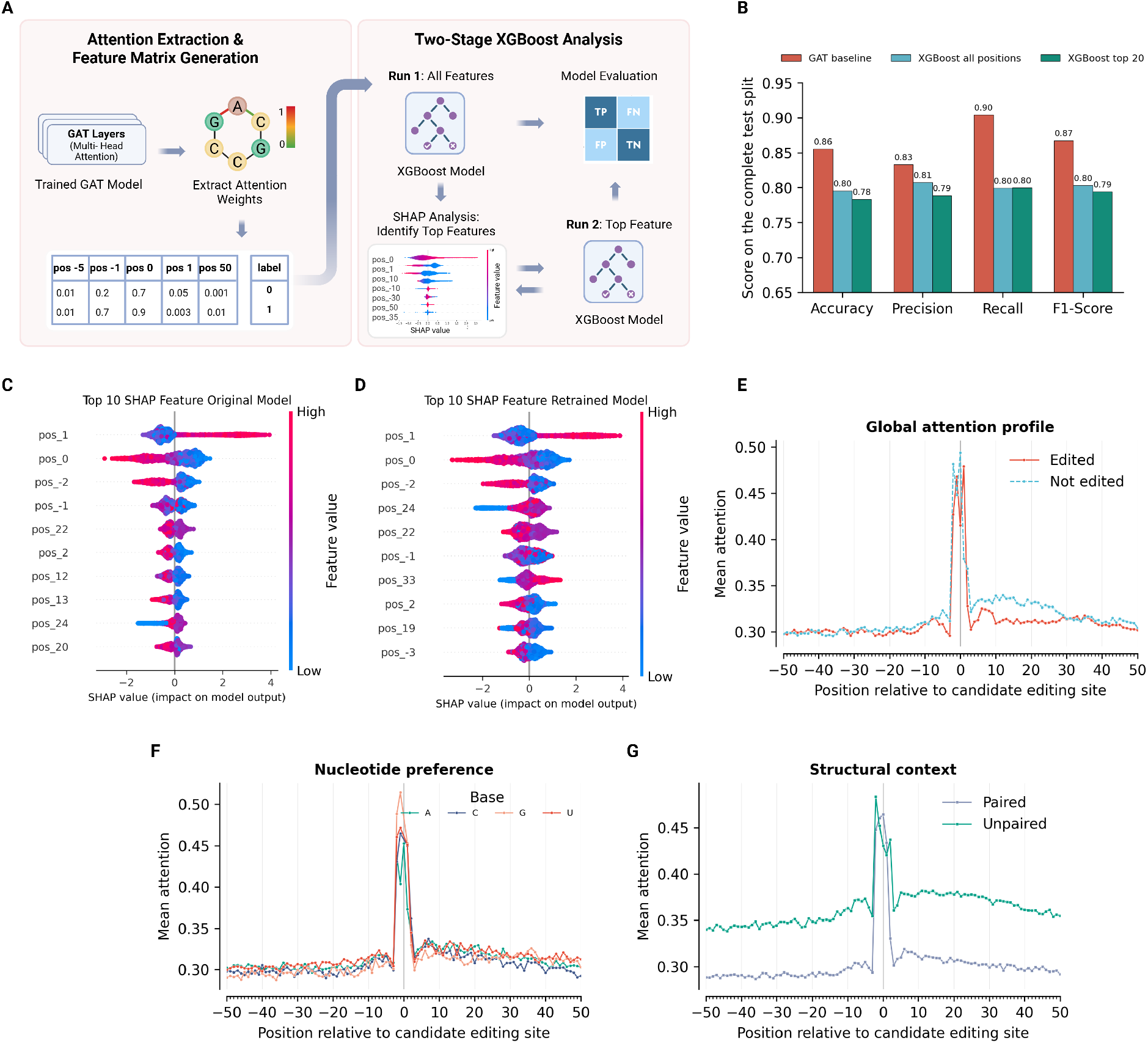
Attention analysis of the baseline model. (A) Interpretability workflow: extraction of edge-level attention weights from the last GAT layer, aggregation into positional features training of an XGBoost classifier on the validation-set attention features, SHAP analysis and selection of the top 20 features on the validation set, and training of a second XGBoost classifier using these features. (B) Held-out test-set Performance metrics (accuracy, precision, recall, F1-score) for three classifiers: baseline GAT (red), XGBoost trained on all attention features (blue), and XGBoost retrained on top 20 SHAP features (green). (C,D) SHAP summary plots showing top 10 attention features for original XGBoost model (C) and retrained model (D), calculated on the validation set, with feature importance values and direction of influence. (E) Mean attention across *±*50 nucleotide windows for edited sites (orange) and non-edited sites (cyan). (F) Mean attention stratified by nucleotide identity (A, U, G, C) at each position relative to the editing site. (G) Mean attention comparing unpaired/loop positions (green) versus base-paired positions (cyan) across the *±*50 nucleotide window Panels E–G include all held-out test sites

The supervised validation demonstrated that attention weights encode substantial predictive information (Figure 5B). Remarkably, an XGBoost classifier trained solely on attention-derived positional features – without access to the original sequence, structure, or any engineered features – achieved strong performance on the Combined dataset: F1=0.80, accuracy=0.80, precision=0.81, and recall=0.80. These robust results demonstrate that attention coefficients alone capture the majority of predictive signal, consistent with the hypothesis that they reflect interpretable positional patterns rather than serving as computational artifacts. As expected, the full baseline model achieved higher performance (F1=0.87, accuracy=0.86), as it leverages additional information from multi-layer graph representations, node embeddings after message passing, and global pooling operations. However, the ability of attention weights alone to achieve F1=0.80 is consistent with the hypothesis that the model’s decision-making process reflects interpretable positional patterns. SHAP analysis of the XGBoost model (Figure 5C) revealed that positions immediately surrounding the editing site (positions 0, *±*1, *−*2) ranked as the most influential features, aligning with experimental findings identifying these nucleotides as critical determinants of editing efficiency. When retrained using only the top 20 SHAP-identified features, the XGBoost model maintained nearly identical performance (F1=0.79, accuracy=0.78; Figure 5B), with SHAP values again highlighting the same proximal positions (Figure 5D), confirming that a small, interpretable subset of positional attention features retains full attention-based predictive power and facilitates biological interpretation. Both XGBoost classifiers were trained on 3,793 validation sites from 139 *Alu* duplexes, with SHAP ranking and top-20 feature selection also confined to the validation split. Final performance was evaluated on 4,864 test sites from 179 duplexes, with no duplex overlap between the validation and test sets; the test data were not used for XGBoost training or feature selection.

Descriptive analysis of attention patterns revealed that the baseline model’s learned representations are consistent with established biochemical rules. Global attention profiles (Figure 5E) showed a sharp attention peak around the candidate editing site (positions *−*2 to +1) for both edited sites (orange line) and non-edited sites (cyan line), with non-edited sites exhibiting slightly elevated attention across much of the downstream window. Stratifying attention by nucleotide identity (Figure 5F) revealed a striking pattern: at the immediate 5’ neighbour (position *−*1), mean attention was highest for guanosine, followed by uracil and cytosine, and lowest for adenosine. Guanosine and uracil represent two particularly informative identities under the established 5’ nearest-neighbour rule – G the strongest suppressor of editing and U the most preferred neighbour – indicating that the model autonomously learned to weight this position by its known influence on ADAR activity. Structural context analysis (Figure 5G) demonstrated that unpaired/loop positions (green line) receive markedly higher mean attention than base-paired positions (blue line) across most of the window, with both structural contexts exhibiting pronounced attention peaks near the candidate editing site.

#### In silico mutagenesis reveals sequence and structure preferences at core editing positions

To systematically probe the baseline model’s learned sequence and structural preferences, we performed comprehensive *in silico* mutagenesis analysis on 1,520 high-confidence edited sites (model prediction *>*0.7 on true edited sites from the validation set). For each edited site, we generated all possible single-nucleotide variants at positions spanning from-3 to +3 relative to the editing site (position 0), maintaining the target adenosine unchanged. At each position, we substituted the original nucleotide with each of the four bases (A, G, C, T/U), updated the corresponding node features in the graph representation, and measured the change in model prediction. To isolate sequence effects from structural effects, we performed two parallel analyses: (1) sequence-only mutagenesis, where only the base identity was altered while preserving the original pairing status, and (2) structure-coupled mutagenesis, where mutations that would disrupt canonical or wobble base pairs (A–U/T, G–C, G–U/T) also triggered structural changes by modifying the paired/unpaired status and removing the corresponding base-pairing edge from the graph. We aggregated results across analysed edited sites to generate mean preference scores for each base at each position, normalized by subtracting the position-wise mean to highlight relative preferences (Figure 6A).

**Figure 6:**
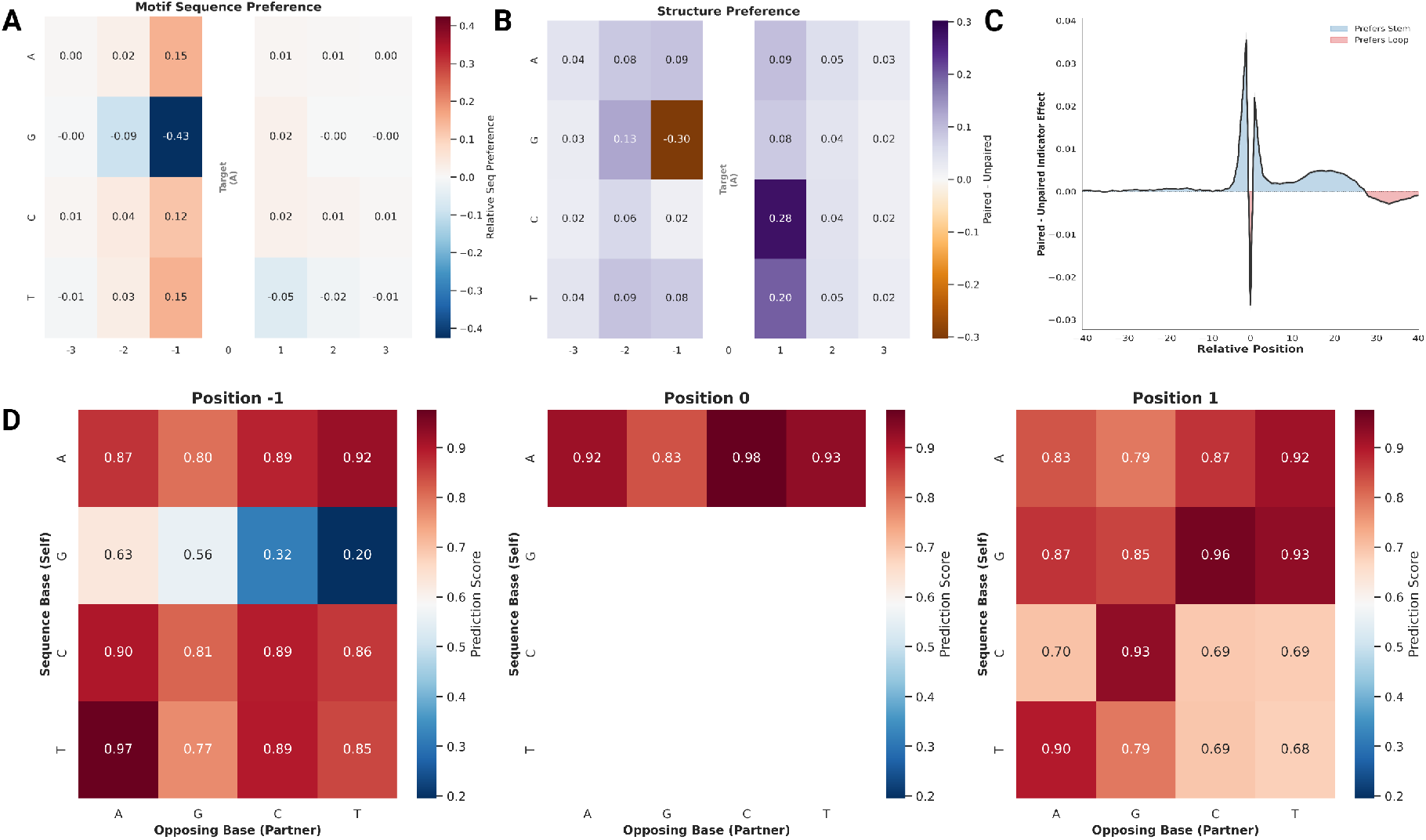
In silico mutagenesis of baseline model reveals sequence and structural preferences. (A) Sequence preference heatmap for positions-3 to +3. Rows show bases A, G, C, U/T; values are normalized prediction scores (position-wise mean subtracted). Red/orange indicates enhanced editing; blue indicates suppression. Position 0 (target A, marked by vertical line) excluded from mutations. (B) Structure-coupled mutagenesis at positions with an existing RNAfold-predicted partner. Values show the mean prediction difference between retaining and disrupting the existing base pair for each substituted base and position. Purple indicates higher predictions when the pair is retained; orange indicates higher predictions when it is disrupted. (C) Sensitivity to the paired-state indicator across positions-40 to +40. At each position, only the focal node’s paired-state indicator was toggled, while sequence identity, graph edges, and all other features were held fixed. The line shows the mean prediction difference between the paired and unpaired indicator states; blue shading indicates positions where the model assigns higher predictions to the paired-state indicator (stem preference), whereas red shading indicates higher predictions to the unpaired-state indicator (loop preference). (D) Base-pairing interaction matrices at positions-1, 0, and +1. Each 4 *×* 4 heatmap shows prediction scores for all combinations of sequence base (rows) and opposing partner base (columns). Higher values (dark red) indicate higher model predictions. Position 0 shows only the A row because the target nucleotide was maintained as adenosine.

**Figure 7:**
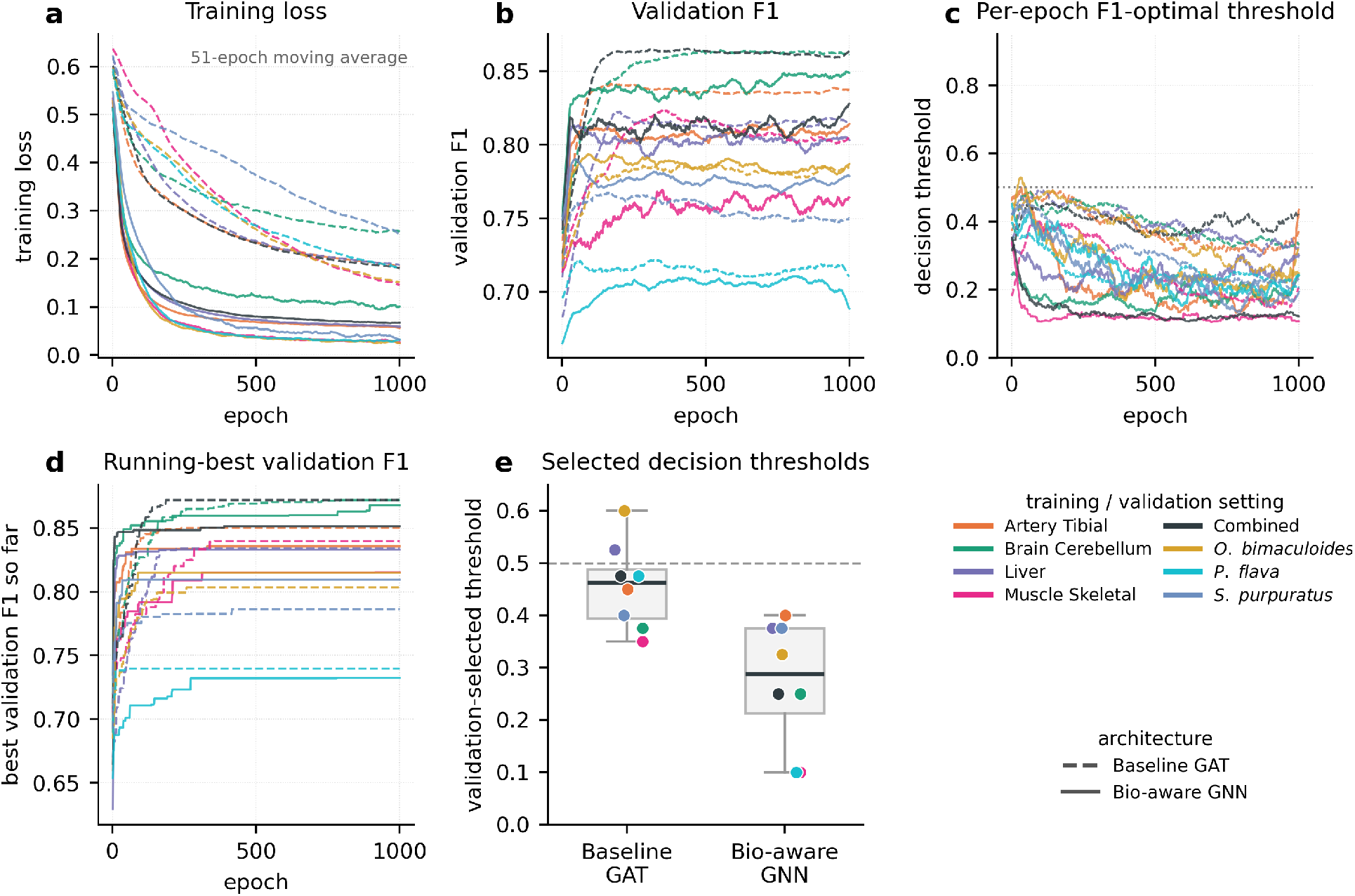
Training dynamics and selected decision thresholds for both architectures. Each colored line is one train*→*(own-tissue/species)-validation setting (Baseline GAT dashed, Bio-aware GNN solid); curves in (a)–(c) are smoothed with a moving-average window of 51 epochs. (a) Training loss over epochs. (b) Validation F1, evaluated at the per-epoch optimal threshold, over epochs. (c) Per-epoch optimal decision threshold (the threshold maximising validation F1 at each epoch; dotted line marks 0.5). (d) Best validation F1 achieved so far (running maximum). (e) Final validation-selected threshold for each of the eight human and within-species settings (four tissue-specific contexts, the GTEx-wide Combined context, and three within-species settings), shown separately for the Baseline GAT and Bio-aware GNN; each point is colored by its training/validation setting. Boxes show the median and interquartile range and the dashed line marks 0.5.

The sequence preference analysis (Figure 6A) revealed striking patterns consistent with established ADAR biochemistry. At position-1 (5’ neighbor), G produced substantially reduced predictions, whereas A, C, and T produced similarly higher predictions, recapitulating the well-documented 5’ neighbor preference rule where upstream purines – particularly G – strongly suppress editing Polson & Bass (1994); Eggington et al. (2011). This suppression extended to position-2, where G also showed reduced preference compared to pyrimidines. At position +1 (3’ neighbor), the effects were smaller: T reduced the prediction, whereas A, C, and G produced modest positive shifts. At the editing site itself (position 0), only adenosine is biologically relevant, so this position was excluded from base-substitution analysis.

The structure-coupled mutagenesis analysis (Figure 6B) revealed distinct pairing preferences that vary by both position and nucleotide identity. At position-1, the model exhibited a striking base-specific structural preference: when guanosine occupies this position, the model strongly prefers it to remain unpaired (positive values in heatmap indicate preference for paired status; negative values indicate preference for unpaired status), consistent with structural studies showing that unpaired G at-1 facilitates ADAR binding Doherty et al. (2022). In contrast, when position-1 contains U, C, or A, the model showed a moderate preference for these bases to be paired, suggesting that structural context requirements differ based on local sequence composition. At position +1, pyrimidines (U and C) exhibited strong preferences to be base-paired, potentially reflecting a requirement for structural stability in the 3’ stem region to properly position the editing site for catalysis. These position-and base-specific structural preferences demonstrate that the model has learned coupled sequence-structure rules rather than treating structure as a position-independent feature.

To examine pairing-state preferences beyond the immediate editing motif, we performed a positional paired-state indicator analysis across a broad window spanning positions-40 to +40 (Figure 6C). For each position, we compared model predictions after setting only the focal node’s paired-state indicator to paired or unpaired, while retaining the graph edges, nucleotide identity, and all other features, and measured the impact as Paired minus Unpaired. Positive values indicate positions where the paired-state indicator increases the model prediction, while negative values indicate positions where the unpaired-state indicator is preferred. The analysis revealed several striking features (Figure 6C). First, position 0 (the editing site) showed the strongest preference for the unpaired-state indicator, consistent with the fundamental requirement for adenosine accessibility. Second, positions *∼*30–35 downstream of the editing site exhibited a consistent, although weaker, preference for the unpaired-state indicator, aligning with previous studies Zambrano-Mila et al. (2023). Third, positions around +20 showed a modest preference for the paired-state indicator, potentially reflecting a contribution of pairing-state information from the mid-3’ region. Fourth, the overall signal was more extended in the downstream (3’) region compared to upstream (5’), suggesting that the model uses pairing-state information asymmetrically around the target. This asymmetry may reflect the directional nature of ADAR substrate recognition. These positional effects indicate that the model’s predictions are sensitive to pairing-state information beyond the immediate local sequence context.

To directly assess whether the model learned canonical base-pairing rules and their influence on editing, we performed a base-pairing interaction analysis at the three core positions (-1, 0, +1) (Figure 6D). For sites with an existing RNAfold-predicted partner at the queried position, we systematically varied both the sequence-side nucleotide and its opposing partner across all 16 possible combinations at positions-1 and +1; at position 0, the target nucleotide was maintained as A while its partner was varied across the four bases. Valid pairs (Watson-Crick: A–U/T, G–C; wobble: G–U/T) were marked as paired (feature value = 1) with the base-pairing edge retained in the graph, while invalid pairs triggered structural disruption: the pairing status was set to 0 and the base-pairing edge was removed from the graph to reflect the absence of stable hydrogen bonding. This topology-aware perturbation allowed us to test whether the model responds appropriately to both sequence changes and their structural consequences. Results are presented as 4 *×* 4 heatmaps where rows represent the sequence-side base and columns represent the opposing partner base, with prediction scores indicating editing probability (Figure 6D).

At position 0 (the editing site), the strongest prediction occurred when adenosine (A) was opposed by cytosine (C), recapitulating the well-established A:C mismatch preference of ADAR enzymes Wong et al. (2001). At position-1, focal G produced lower predictions across all partner identities than focal A, C, or U/T. Within the G row, however, the G:A and G:G mismatches produced higher predictions than G:C and G:U/T, consistent with previous studies showing that a G:G mismatch at-1 can create a local structural distortion that facilitates ADAR binding Doherty et al. (2022). Conversely, when position-1 contained uracil (U/T) with guanosine (G) opposite, predictions were substantially reduced, matching experimental findings that U:G wobble pairs at-1 suppress editing and can be exploited for site-directed RNA editing control Reautschnig et al. (2025). More broadly, position +1 showed a general preference for stable base-pairing (strong signals along the Watson-Crick diagonal: A:T, G:C), reflecting the requirement for structural integrity in the 3’ stem to position the editing site correctly. Together, these interaction analyses demonstrate that the baseline model autonomously learned not only individual base preferences but also context-dependent base-pairing rules, consistent with the hypothesis that graph-based structural encoding captures the coupled sequence-structure logic governing ADAR substrate recognition.

## 3 DISCUSSION

ADAREDIT demonstrates that integrating RNA secondary structure explicitly into predictive models of A-to-I editing supports both strong predictive accuracy and biological interpretability. Unlike sequence-based or structure-implicit approaches, the baseline graph model captures spatial patterns associated with ADAR substrate recognition – such as position-specific pairing, local asymmetry, and inhibitory neighbor bases – without requiring manually encoded biochemical features. Evaluated on held-out test sets in which each *Alu* pair and its predicted structure appears in a single split, the model attains strong within-and cross-tissue performance. Editing levels at sites co-measured in an independent human RNA-seq cohort, RNAAtlas, were highly concordant with the GTEx-derived measurements (Supplementary Fig. S3), supporting their robustness across cohorts and processing pipelines. The compact baseline graph and the biochemically enriched bio-aware variant perform comparably in this regime; we therefore view the structure-native graph formulation itself – the explicit encoding of base-pairing topology – as the central methodological advance, with the two variants serving as complementary members of a single ADAREDIT family.

A single model trained jointly over all five tissue contexts, with one shared encoder and a context-specific output head, retained full context-specific resolution while reducing the five-model setup to a single predictor. The joint baseline performed similarly to the separate per-tissue baselines, whereas the joint bio-aware model achieved modestly higher average F1 and AUROC than the corresponding separate bio-aware models and slightly outperformed the joint baseline on average, although the differences varied across tissue contexts. Thus, the results suggest that the bio-aware encoder benefits modestly from the higher-supervision joint setting: although the two encoders were comparable when each tissue was modeled alone, the richer biochemical annotations may become more useful when the shared encoder receives labels from multiple tissue contexts for the same underlying sequence–structure substrates. Performance remained comparable when the Combined label was excluded from joint-model supervision, indicating that the joint-model results were not dependent on this partially redundant label (Supplementary Fig. S8). Because the present benchmark is dominated by stable inverted *Alu* hairpins, this result should be interpreted within the *Alu*-derived regime rather than as proof of universal dsRNA editability modeling.

Beyond predictive performance, ADAREDIT may offer future value for the design of site-directed RNA editing therapies. Recent advances have shown that endogenous ADAR enzymes can be co-opted using synthetic guide RNAs to correct pathogenic mutations. However, efficient guide design remains challenging, often requiring laborious empirical screening. ADAREDIT could potentially address this bottleneck by enabling computational prioritization of candidate guides across the transcriptome. Rather than testing dozens or hundreds of guide configurations in vitro, researchers can use the model to rapidly score large design spaces and focus experimental validation on the most promising candidates – reducing cost, time, and design uncertainty. In its current form ADAREDIT predicts endogenous editability of naturally occurring *Alu* substrates rather than the efficiency of engineered guide RNA–target duplexes, which differ in dsRNA geometry (shorter stems and mismatches introduced to recruit ADAR); it is therefore best positioned to prioritise candidate adenosines for approaches that recruit or leverage endogenous ADAR1 activity. This ADAR1-oriented scope remains therapeutically relevant because ADAR1 is broadly expressed across tissues and is a practical endogenous effector for guide-directed editing in accessible peripheral contexts, including liver. Nevertheless, applying ADAREDIT directly to guide design would require additional validation on guide RNA–target pairs, and direct guide-RNA design and broad recoding of ADAR2-dominant targets remain future extensions rather than validated current applications.

While the model retains predictive signal across the evaluated tissue and species settings, its performance on certain ADAR2-dominant targets remains modest. This likely reflects the composition of the training data: ADAREDIT was trained primarily on *Alu* elements, which are abundant in the human transcriptome and predominantly edited by ADAR1. Expanding the training set to include a broader repertoire of coding-region edits and validated ADAR2 substrates will likely improve the model’s sensitivity to isoform-specific rules. The framework itself is agnostic to ADAR subtype, and can naturally incorporate such data as it becomes available.

Looking ahead, models like ADAREDIT could help shift RNA editing design from empirical trial-and-error toward predictive, computation-guided strategies. With future validation on guide RNA–target pairs, such models could support the prioritization of candidate sites and guide configurations and thereby streamline experimental workflows. As the field progresses, incorporating cell-type context, isoform preferences, and experimentally measured RNA structures may further improve prediction accuracy and broaden clinical applicability.

### Scope and limitations

#### Scope

ADAREDIT is designed for predicting A-to-I RNA editing sites in double-stranded RNA contexts where ADAR enzyme recognition depends on both sequence and secondary structure features. The model operates on computationally predicted RNA secondary structures (RNAfold) representing dsRNA duplexes. It has been trained and validated across five human contexts with diverse ADAR expression profiles (Brain Cerebellum, Artery Tibial, Liver, Muscle Skeletal, Combined) and three evolutionarily distant non-mammalian species lacking *Alu* elements (sea urchin, acorn worm, octopus). The framework is intended for transcriptome-scale prioritization of candidate editing sites and mechanistic interpretation of the structural and sequence determinants governing ADAR specificity. Its primary validated regime is the human *Alu*-derived benchmark evaluated on held-out test sets across the five tissue contexts, supplemented by targeted coding-site scope tests and conservative cross-species transfer analyses. While the graph-based architecture is generalizable in principle to any dsRNA context where secondary structure can be predicted or experimentally determined, the present manuscript supports its strongest claims in the stable, *Alu*-dominated, ADAR1-like endogenous regime represented in training.

#### Limitations

Several technical and biological limitations warrant consideration. First, ADAREDIT relies entirely on computationally predicted secondary structures from RNAfold minimum free energy models, which may not capture dynamic conformational ensembles or tertiary interactions that influence editing. Second, the model does not account for additional RNA-binding proteins that can compete with or facilitate ADAR binding to substrates, thereby modulating editing levels in cellular contexts. Third, binary classification (edited vs. non-edited) does not provide quantitative prediction of editing stoichiometry – a limitation for therapeutic applications requiring precise dosage control – although threshold-relaxation analyses indicate that the within-domain signal is not restricted to a single editing-level cutoff (Supplementary Fig. S1). Fourth, interpretability analyses via attention weights, SHAP, and in silico mutagenesis provide hypothesis-generating signals about which features the model uses for prediction, but they do not establish causality or direct molecular mechanisms. We therefore frame these analyses as consistency checks against known ADAR biochemical preferences rather than as definitive proof of mechanism, and experimental validation is required to confirm the biological relevance of identified patterns and rule out spurious correlations. Fifth, the released human preprocessing path is reference-based rather than donor-personalized; an empirical strand-aware comparison against donor whole-genome sequencing indicates that A-to-G variant contamination affects only a negligible fraction of labeled sites (*∼*0.15%; Supplementary Fig. S4), so this is a bounded rather than a confounding factor. Finally, the model has been evaluated primarily within the *Alu*-derived duplex paradigm, so that recoding/ADAR2 transfer, synthetic guide-RNA design, and the cross-species benchmark remain scope-limited – coding-target sample sizes are small and the non-*Alu* species negatives rely on a proximity-based labeling rule (Supplementary Fig. S5); broader validation across diverse non-*Alu* substrates, alternative splicing isoforms, and species-specific editing contexts remains an important direction for future development.

## Data Availability

The processed, model-ready datasets and fixed train, validation, and test partitions used in this study are publicly available at https://github.com/Scientific-Computing-Lab/AdarEdit. Human editing data were derived from GTEx v8 RNA-seq measurements across 47 tissues. RNAAtlas RNA-seq data used for the independent-cohort concordance analysis are available under NCBI BioProject accession PRJNA576920. Cross-species editing sites were obtained from Zhang et al. Zhang et al. (2023). The repository contains the processed data underlying the reported model evaluations and analyses.

## Code Availability

The source code used for data processing, model training and evaluation, and generation of the reported analyses and figures is publicly available at https://github.com/Scientific-Computing-Lab/AdarEdit. The repository also provides the trained model checkpoints, documented software environments, and scripts for reproducing the baseline and bio-aware models, cross-tissue and cross-species evaluations, attention and SHAP analyses, component ablations, and *in silico* mutagenesis experiments.

## LLM Usage

During preparation of this manuscript and its associated reproducibility materials, the authors used generative-AI tools to assist with manuscript drafting and language editing, code review and debugging, reproducibility auditing, and the preparation and documentation of analysis and visualization scripts. The authors take full responsibility for the content of the manuscript and the associated code.

## Acknowledgments

This work was supported by US National Institutes of Health award R35GM122543 (M.L.), Foundation Fighting Blindness (grants TA-GT-0620-0790-HUJ and PPA-0923-0865-HUJ), grants from the Israeli Ministry of Science (grant 3-17916), and by Israel Science Foundation (2637/2). M.L. is the Robert W. and Vivian K. Cahill Professor of Cancer Research. E.Y.L. is a fellow at the Israel Institute of Advanced Studies.

## **A** Previous Work on A-to-I RNA Editing Prediction

Computational prediction of A-to-I editing sites has progressed through several methodological phases, reflecting a steady shift from manually designed descriptors to representation learning. Early predictors relied on classical machine learning models (e.g., support vector machines, random forests, and XGBoost) built on handcrafted features such as local sequence motifs, nucleotide composition around candidate sites, thermodynamic properties derived from RNA secondary structure, and evolutionary conservation scores Tac et al. (2021); Ouyang et al. (2018); Liu et al. (2021); Jiang et al. (2024); Chen et al. (2016). Although these approaches achieved moderate accuracy, their reliance on predefined biological descriptors constrained the discovery of editing determinants outside the chosen feature set and often required substantial manual re-engineering when transferred across datasets or conditions. Recent work continues to explore combinations of sequence and secondary-structure descriptors in broader evaluation settings, including cross-testing and conservation-oriented analyses Zawisza-Á lvarez et al. (2024).

With the rise of deep learning, convolutional and recurrent architectures began to learn predictive patterns directly from sequence windows, reducing dependence on explicit feature engineering. EditPredict Wang et al. (2021); Ouyang et al. (2018) uses convolutional networks on 201-nucleotide windows to classify editing sites, while other frameworks combine multiple sequence-derived views or extend prediction to alternative sequencing modalities (e.g., nanopore) Chen et al. (2023b,a). More recently, PreAIS Fang et al. (2025) further illustrates the continued strength of deep learning for direct site prediction using DNN/CNN-based modeling and modern interpretability analyses. Despite these advances, most deep learning predictors still treat RNA primarily as a linear string, and structural context is typically incorporated only indirectly (as derived features) rather than encoded as an explicit relational substrate.

Large pre-trained models have also been adapted to editing prediction, motivated by their ability to capture richer contextual dependencies. RNA-FM Chen et al. (2022), trained on millions of RNA sequences with a masked-language objective, can be fine-tuned for downstream editing tasks. ADAR-GPT Rosenwasser et al. (2026) applies continual fine-tuning of GPT-4o-mini on sequence windows and shows competitive performance when benchmarked against established deep learning baselines. In a related therapeutic setting, Helix Cao et al. (2025) augments transformer attention with base-pairing probability matrices to predict on-target editing efficiency for guide RNA design, reporting Spearman correlations of approximately 0.84 between predicted and measured on-target efficiency.

Helix and ADAREDIT target complementary problems. Helix predicts the on-target efficiency of engineered guide RNAs – a continuous, guide-optimisation objective evaluated by Spearman correlation on designed guide–target duplexes – whereas ADAREDIT predicts the endogenous editability of naturally occurring *Alu* adenosines, a binary classification evaluated by F1 and AUROC. Because the two operate on different substrates (engineered guide–target pairs versus endogenous inverted-*Alu* duplexes) and quantify different outcomes (editing efficiency versus edited/non-edited status), their reported scores measure different quantities and do not share a common scale. Their structural representations also differ: Helix supplies base-pairing probabilities as soft features to a transformer over the linear sequence, whereas ADAREDIT represents the duplex explicitly as a graph of backbone and typed base-pair edges, enabling message passing over the secondary-structure topology itself.

In parallel, complementary lines of work focus on improving the reliability of editing labels and calling editing events directly from RNA-seq, including deep learning frameworks designed to separate RNA editing from DNA variation Fu et al. (2024) and temporal convolutional classifiers trained on large collections of known A-to-I events Fonzino et al. (2025). These efforts are valuable for curating high-confidence training sets, but they do not directly address the core representational challenge of encoding dsRNA geometry as a native relational structure for site-level prediction.

Graph neural networks have been widely used to model RNA structure and function in adjacent domains, including RNA secondary structure learning Zhang et al. (2021), RNA–protein interaction prediction Yamada & Hamada (2022); Saon et al. (2024), and hybrid architectures that integrate language model embeddings with graph attention for binding-site tasks Xiao et al. (2024). Structure-aware non-GNN predictors have also incorporated RNAfold-derived base-pairing information into editing classification Liu et al. (2021). However, A-to-I editing-site predictors remain overwhelmingly sequence-centric – CNNs, RNNs, transformers, or feature-based models – rather than graph-native architectures that explicitly encode both sequential adjacency and base-pairing relationships within a unified representation. We are not aware of prior graph-based predictors purpose-built for A-to-I editing-site classification that natively integrate these edge types while providing attention-level interpretability aligned with established ADAR biochemistry. This positions ADAREDIT in a distinct methodological space relative to existing editing predictors and the broader GNN-for-RNA literature.

## **B** Comprehensive Performance Metrics Across All Experimental Settings

Table 1 presents performance metrics across 36 evaluation settings: 25 train*→*test combinations across four tissue-specific human contexts and the GTEx-wide Combined context, six cross-species evaluations involving three evolutionarily distant non-*Alu* organisms, and five context-specific outputs from the joint models.

For each setting, we report dataset sizes and, for both the bio-aware and baseline encoders, F1, precision, recall, AUROC, and AUPRC at the validation-selected decision threshold, which is also tabulated. Across the four tissue-specific within-context evaluations, F1 differed by less than 0.01 between the two encoders; in the Combined context, the baseline was higher by 0.032. Across the full matrices, the baseline attained the higher F1 in 19 of 25 human train*→*test combinations and four of six cross-species settings, consistent with the compact baseline graph capturing much of the editability signal while the bio-aware variant provides comparable, complementary performance. Among the two joint models, the joint bio-aware encoder achieved higher AUROC than the joint baseline in all five output contexts and higher F1 in Artery, Brain, Liver, and Muscle Skeletal, whereas the joint baseline was slightly higher in F1 for Combined. Relative to the corresponding independently trained bio-aware models, joint training improved both F1 and AUROC in Artery, Combined, Liver, and Muscle Skeletal, while Brain remained effectively unchanged. These results indicate a modest, context-dependent benefit from pooled training, particularly for the bio-aware encoder.

## **C** Training Dynamics and Checkpoint Robustness

We characterized the training behavior of both architectures across the 16 per-tissue and within-species training runs shown in Figure 7. These analyses address the concern that F1-optimized checkpoint selection might reflect isolated high-performing epochs rather than stable training behavior. The smoothed training-loss trajectories (Figure 7a) decline substantially over training and approach stable low-loss regimes, with no evidence of late-epoch divergence. Validation F1 (Figure 7b) rises rapidly and then fluctuates around sustained high-performance regimes, while the running-best validation F1 (Figure 7d) increases gradually and is maintained once reached. Thus, the selected check-points occur within stable training regimes rather than reflecting numerical instability or failed convergence.

The per-epoch F1-optimal decision threshold (Figure 7c) is generally below 0.5 but can fluctuate across epochs because nearby thresholds often yield similar validation F1. The final validation-selected thresholds (Figure 7e) show the same calibration pattern: most lie below 0.5, with the Baseline GAT at a higher median threshold than the Bio-aware GNN (0.463 versus 0.288). These thresholds determine only the operating point for threshold-dependent metrics such as F1; the AUROC and AUPRC values reported throughout are threshold-free. Overall, the training logs support the interpretation that performance is driven by stable validation behavior across the per-tissue and within-species runs, rather than by isolated checkpoint or threshold artifacts.

## Supplementary Information

Editing-level distribution and threshold relaxation

**Figure S1:**
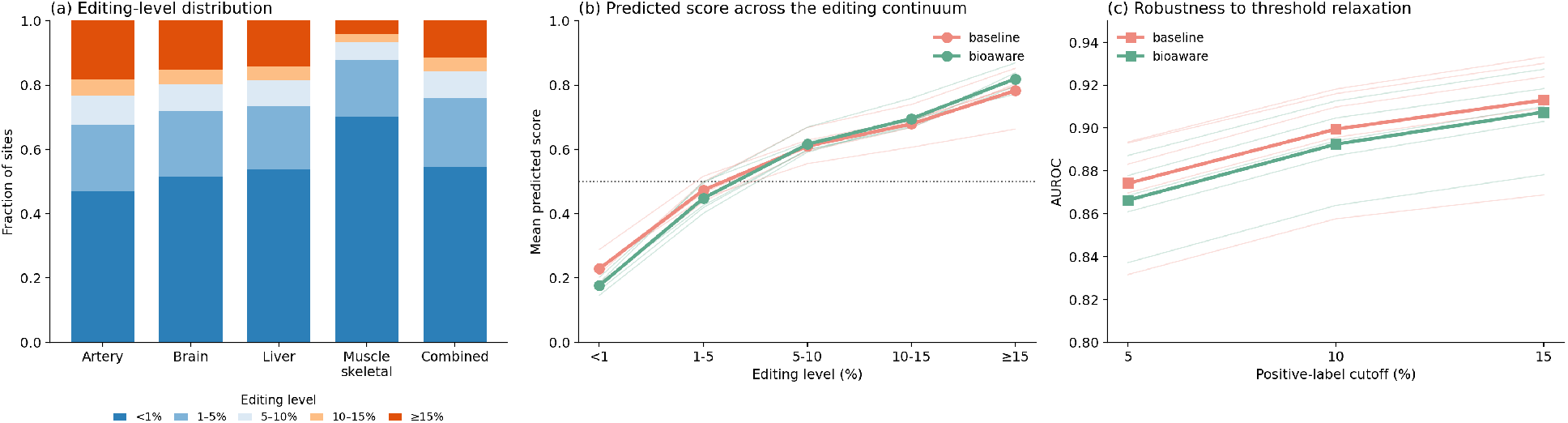
Intermediate editing sites and robustness to threshold relaxation. (a) Distribution of editing levels across all GTEx sites within *Alu* pairs prior to binary filtering, stacked by editing-level bin and shown per context. Intermediate sites (1–15%) constitute 25.7–34.9% of the data across the five contexts. (b) Mean predicted score of the Baseline GAT and Bio-aware GNN as a function of the true editing level, for every held-out intermediate (1–15%) site together with the binary extremes. Although the models are trained only on the *<*1%/*≥*15% extremes, the predicted score rises monotonically with editing level (faint lines, individual tissues; bold lines, mean across tissues). (c) AUROC on the held-out test sets when the positive-label cutoff is relaxed to *≥*5%, *≥*10%, and *≥*15% (negatives fixed at *<*1%, the intermediate (1%, cutoff) zone excluded). Discrimination remains high (AUROC *≥* 0.83 in every tissue) even at the most lenient cutoff. Full per-tissue results are in Table S1.

**Table S1:** Threshold-relaxation results across all five contexts. AUROC for the baseline and bio-aware models at three positive-label cutoffs (*≥*5%, *≥*10%, *≥*15%), scoring all processed coverage-filtered sites on the held-out pair-disjoint test pairs without retraining. AUROC increases monotonically with a stricter positive cutoff in every tissue, and remains above 0.83 even at the most lenient *≥*5% cutoff, indicating that performance is not restricted to the binary training threshold. The *≥*15% column equals the binary held-out test AUROC reported in the main text.

| Context | ≥5% |  | ≥10% |  | ≥15% |  |
| --- | --- | --- | --- | --- | --- | --- |
|  | Baseline | Bio-aware | Baseline | Bio-aware | Baseline | Bio-aware |
| Artery Tibial | 0.883 | 0.878 | 0.910 | 0.905 | 0.924 | 0.918 |
| Brain Cerebellum | 0.893 | 0.887 | 0.916 | 0.913 | 0.930 | 0.927 |
| Liver | 0.870 | 0.869 | 0.896 | 0.894 | 0.909 | 0.910 |
| Muscle Skeletal | 0.832 | 0.837 | 0.858 | 0.864 | 0.869 | 0.878 |
| Combined | 0.894 | 0.861 | 0.918 | 0.887 | 0.933 | 0.903 |

**GTEx tissue-selection rationale**

**Figure S2:**
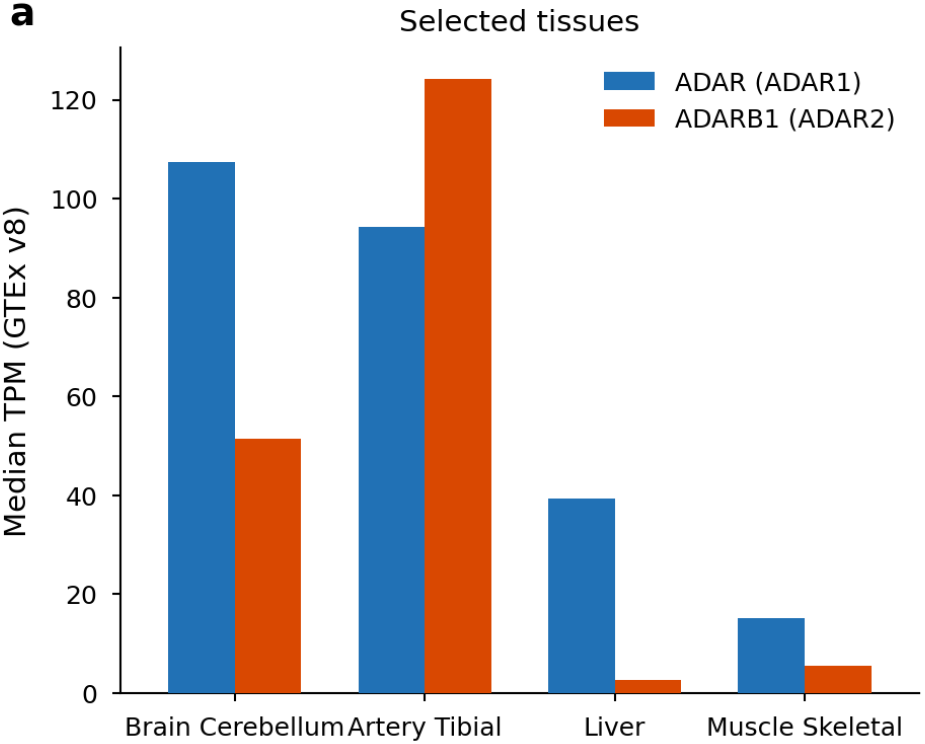
ADAR1 and ADAR2 expression across the selected tissues. Median GTEx v8 expression (TPM) of *ADAR* (ADAR1) and *ADARB1* (ADAR2) for the four individual tissues (the Combined dataset pools editing sites across all 47 GTEx v8 tissues and therefore has no single expression profile). Liver is ADAR1-dominant with minimal ADAR2 (*ADAR* 39.4, *ADARB1* 2.6); Artery Tibial has the highest *ADARB1* of any GTEx v8 tissue (124.3, exceeding *ADAR* 94.3); Brain Cerebellum expresses both isoforms highly (*ADAR* 107.4, *ADARB1* 51.4), additionally expresses the brain-restricted *ADAR3*, and shows the highest *Alu* editing index across tissues; Muscle Skeletal has the lowest ADAR1 (15.3) and the lowest editing index, providing a low-signal stress test.

**Independent human cohort: RNAAtlas re-measurement of *Alu* editing**

**Figure S3:**
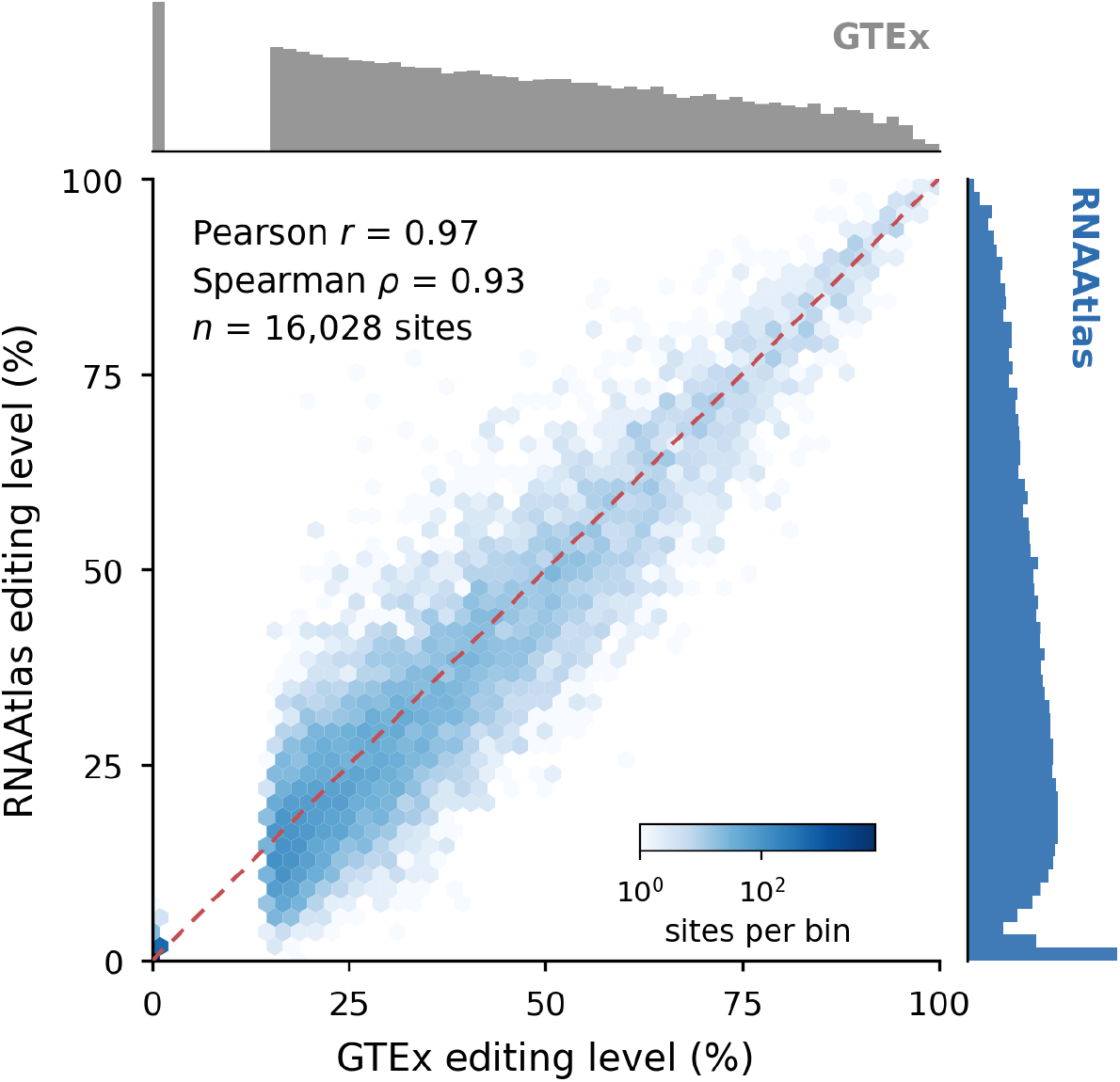
Independent re-measurement of *Alu* editing in the RNAAtlas cohort. Editing levels for *Alu* adenosines were quantified de novo from RNAAtlas (PRJNA576920; 216 samples) and compared with the GTEx editing levels used here. Central panel, per-site editing level in the two cohorts (*n* = 16,028 co-measured sites; median RNAAtlas coverage 450 reads; dashed line, identity); marginal histograms, the editing-level distribution in each cohort. Sites were classified as edited (*≥*15%) or non-edited (*<*1%) on GTEx levels; the independent RNAAtlas re-measurement is continuous. Of these sites, 1,678 fell within the intermediate RNAAtlas range (1–15%); none crossed directly from the GTEx non-edited to the RNAAtlas edited range, and one crossed from the GTEx edited to the RNAAtlas non-edited range (0.006%).

**Donor-versus-reference SNP audit**

**Figure S4:**
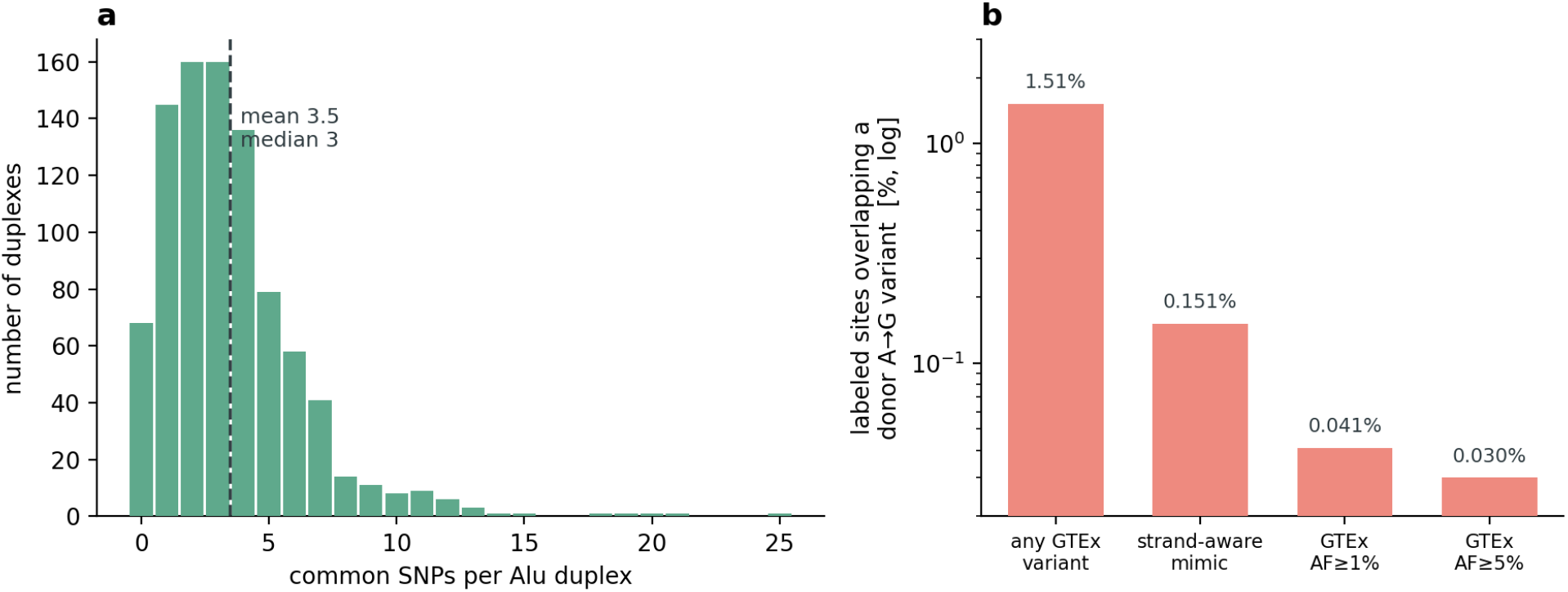
Common population variation is present across the substrates, but donor-level variation at the labelled adenosine is rare and label-balanced. (a) Distribution of common germline SNPs (UCSC Common SNPs 150; minor-allele frequency *≥*1% in at least one population) per *Alu*-pair duplex across the 905 initially reconstructed candidate duplexes. Each duplex (*∼*586 nt) carries on average 3.5 common SNPs (median 3); 92.5% carry at least one, corresponding to *∼*0.59% of positions. (b) Overlap of the 100,377 labelled sites with the actual donor genotypes from GTEx v8 whole-genome sequencing (838 donors, SHAPEIT2-phased), at increasing stringency: any donor variant at the exact coordinate (1.51%); a strand-aware editing-mimicking variant–an A*→*G substitution on the transcribed strand, i.e. genomic A*→*G for + strand and T*→*C for *−* strand sites, the only class able to mimic A-to-I editing (0.151%); and donor allele-frequency thresholds (AF *≥*1%, 0.041%; AF *≥*5%, 0.030%). The strand-aware overlap was similar between sites labelled as edited in at least one context (0.175%) and sites labelled as non-edited in all included contexts (0.145%).

**Node and edge feature specification**

**Table S2:** Node and edge feature specification for both architectures. Full formal specification of the node and edge feature vectors for the Baseline GAT and the Bio-aware GNN. **The Baseline GAT carries no edge feature vector (edge dimension** = 0**): backbone and base-pair edges are homogeneous, untyped graph adjacencies and the model receives only topology.** The Bio-aware GNN assigns each edge a discrete type label (0 = sequential backbone, 1 = canonical Watson–Crick, 2 = wobble) and a 12-dimensional continuous attribute vector; internally the type label is embedded (6 dimensions) and concatenated with the continuous attribute vector, then mapped by a small MLP to the 16-dimensional edge attribute consumed by the GATv2 layers. Dimensions were verified directly against the PyTorch Geometric graph cache of the training split.

| (a) Node features |  |  |  |
| --- | --- | --- | --- |
| Model | Feature group | Dim. | Description |
| Baseline GAT | Base identity (one-hot) | 5 | Nucleotide identity A/G/C/U(T)/N |
| Baseline GAT | Paired flag | 1 | 1 if base-paired in the dot-bracket structure |
| Baseline GAT | Relative position | 1 | Signed offset from the queried adenosine |
| Baseline GAT | Target flag | 1 | 1 only at the queried adenosine |
| **Baseline GAT** | **Total** | **8** |  |
| Bio-aware GNN | Core node features | 8 | Base identity, paired flag, target flag, and relative position normalized by sequence length |
| Bio-aware GNN | Left-neighbour identity | 5 | One-hot identity of the 5' neighbour |
| Bio-aware GNN | Right-neighbour identity | 5 | One-hot identity of the 3' neighbour |
| Bio-aware GNN | Stem-loop geometry | 3 | Stem length, loop length, distance to nearest junction |
| Bio-aware GNN | Pair-energy score | 1 | Pairing strength by identity: G–C –3.4, A–U –2.1, G–U –1.0; else 0 |
| **Bio-aware GNN** | **Total** | **22** |  |
| (b) Edge features |  |  |  |
| Model | Edge encoding | Dim. | Description |
| Baseline GAT | Homogeneous edges | 0 | Backbone and base-pair edges identical; untyped, no edge features |
| Bio-aware GNN | Edge type label | 1 | 0 = backbone, 1 = Watson–Crick, 2 = wobble |
| Bio-aware GNN | Continuous edge attributes | 12 | is_seq, is_pair, normalized distance, stem/loop/junction geometry at both ends, pair-energy score, target proximity at both ends |
| **Bio-aware GNN** | **Total** | **13** |  |

**Cross-species negative-distance sensitivity**

**Figure S5:**
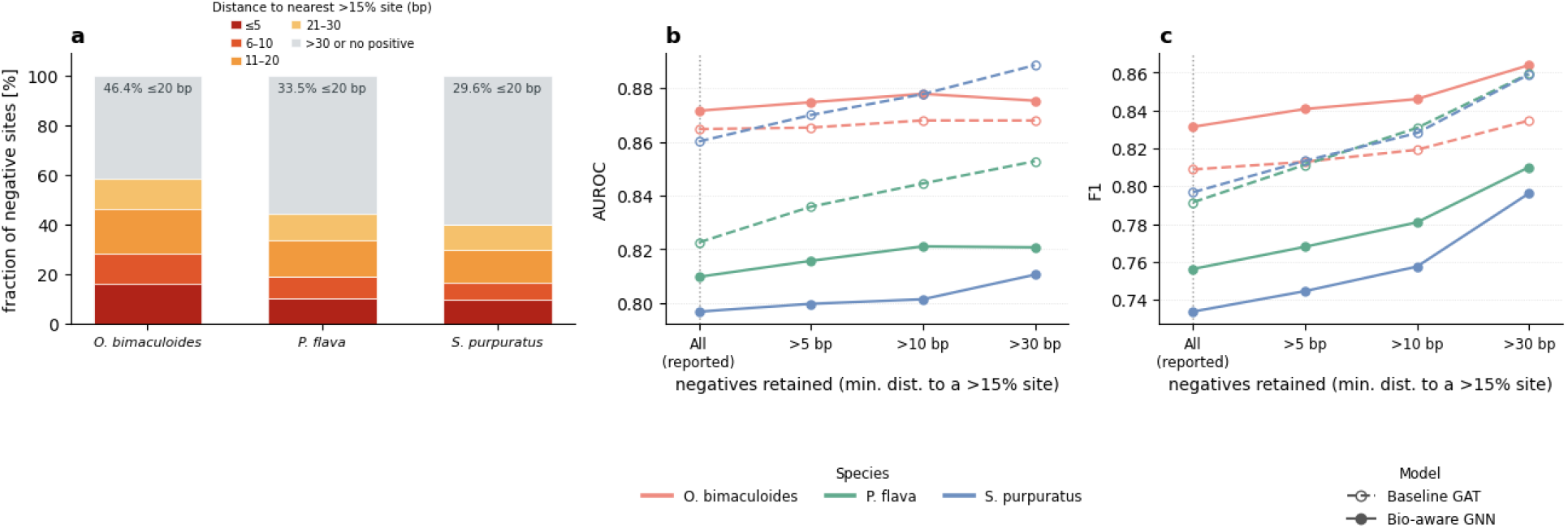
Sensitivity of cross-species performance to the operational negative-labeling rule. (a) Distance from each operational negative (editing level *<* 0.1%; unreported candidate adenosines were encoded as 0) to the nearest benchmark-positive site (*>* 15% editing) in the same dsRNA region and arm. Distances were calculated from the complete filtered and deduplicated datasets before class balancing, comprising 30,655, 13,083, and 27,943 operational negatives in *O. bimaculoides*, *P. flava*, and *S. purpuratus*, respectively. Of these, 46.4%, 33.5%, and 29.6% were within 20 bp of a benchmark-positive site. Negatives without a benchmark-positive site in the same region and arm were retained in the final distance category. Consecutive benchmark-positive sites had a minimum spacing of 1 bp and a median spacing of 4 bp in all three species. (b) AUROC and (c) F1 for the Baseline GAT and Bio-aware GNN on the held-out within-species test sets. Starting from the complete test set, operational negatives within 5, 10, or 30 bp of a benchmark-positive site were progressively excluded. Predictions and validation-selected thresholds were held fixed, and the models were not retrained. F1 increased after filtering, whereas AUROC remained stable or increased modestly, showing that the near-positive negatives do not constitute an artificially easy subset that inflates cross-species discrimination.

**Substrate structural stability**

**Figure S6:**
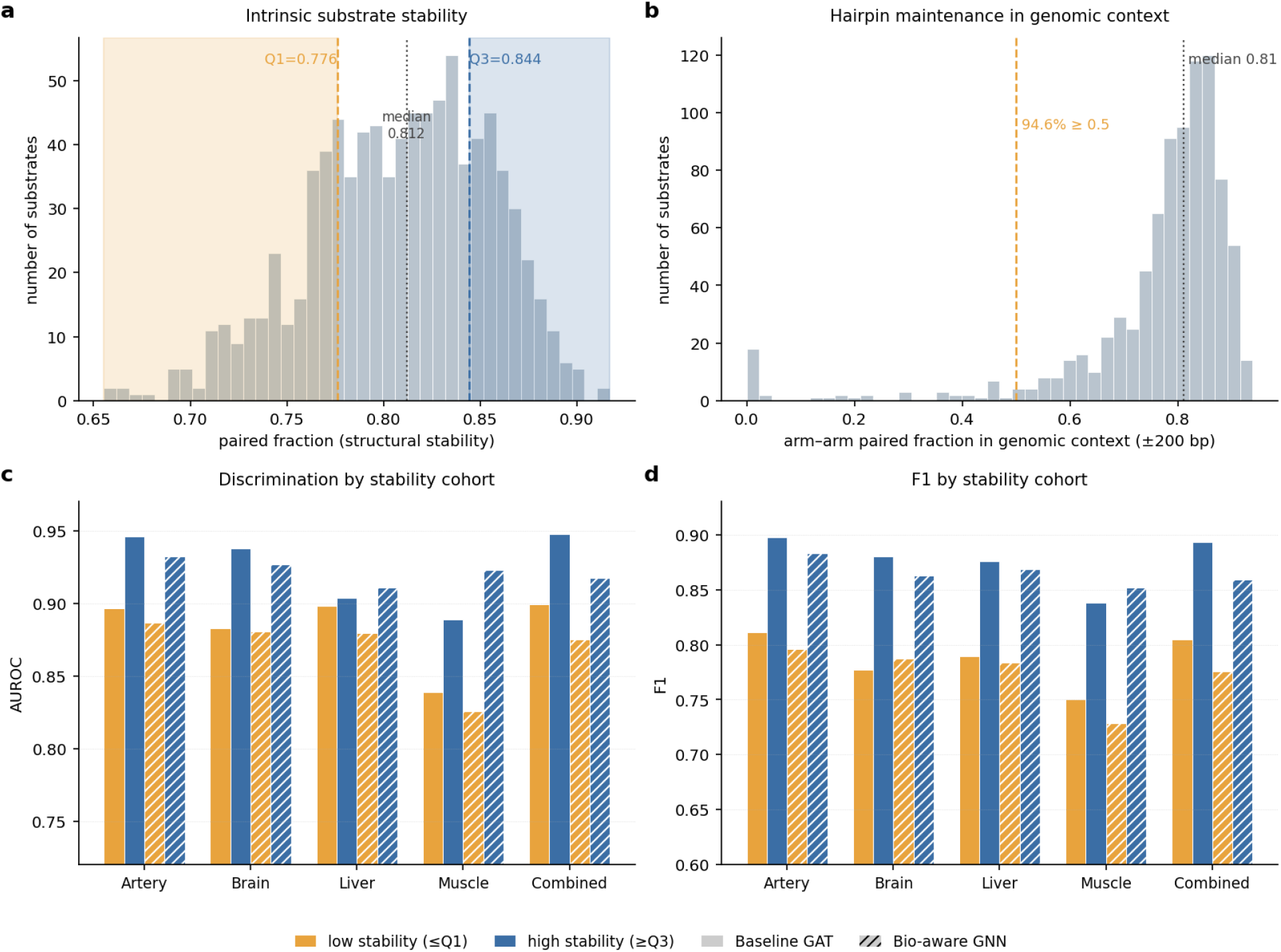
Substrate structural stability and held-out performance across the stability range. (a) Distribution of the paired fraction (fraction of nucleotides engaged in a base pair), used here as a proxy for structural stability, across the input RNAfold structures of the 884 unique *Alu* inverted-repeat substrates (min 0.655, Q1 0.776, median 0.812, Q3 0.844, max 0.917); the dashed lines and shading mark the low-stability (*≤*Q1) and high-stability (*≥*Q3) cohorts. (b) Distribution of the arm-1–arm-2 paired fraction (dsrna frac) when each of 865 pairs is re-folded with *±*200 bp of natural flanking hg38 sequence (no artificial linker; pairs within the folding-length cap); median 0.811, and 94.6% of pairs reach dsrna frac *≥*0.5. (c) Held-out test AUROC and (d) F1 of the Baseline GAT (solid bars) and Bio-aware GNN (hatched bars) on the low-stability (*≤*Q1) and high-stability (*≥*Q3) cohorts, for each of the five human settings (Artery, Brain, Liver, Muscle Skeletal and Combined). Cohorts are defined by the global quartiles in panel a; AUROC and F1 are computed on the pair-disjoint held-out test split, with the validation-selected threshold applied only to F1.

**Component ablation of the bio-aware model**

**Figure S7:**
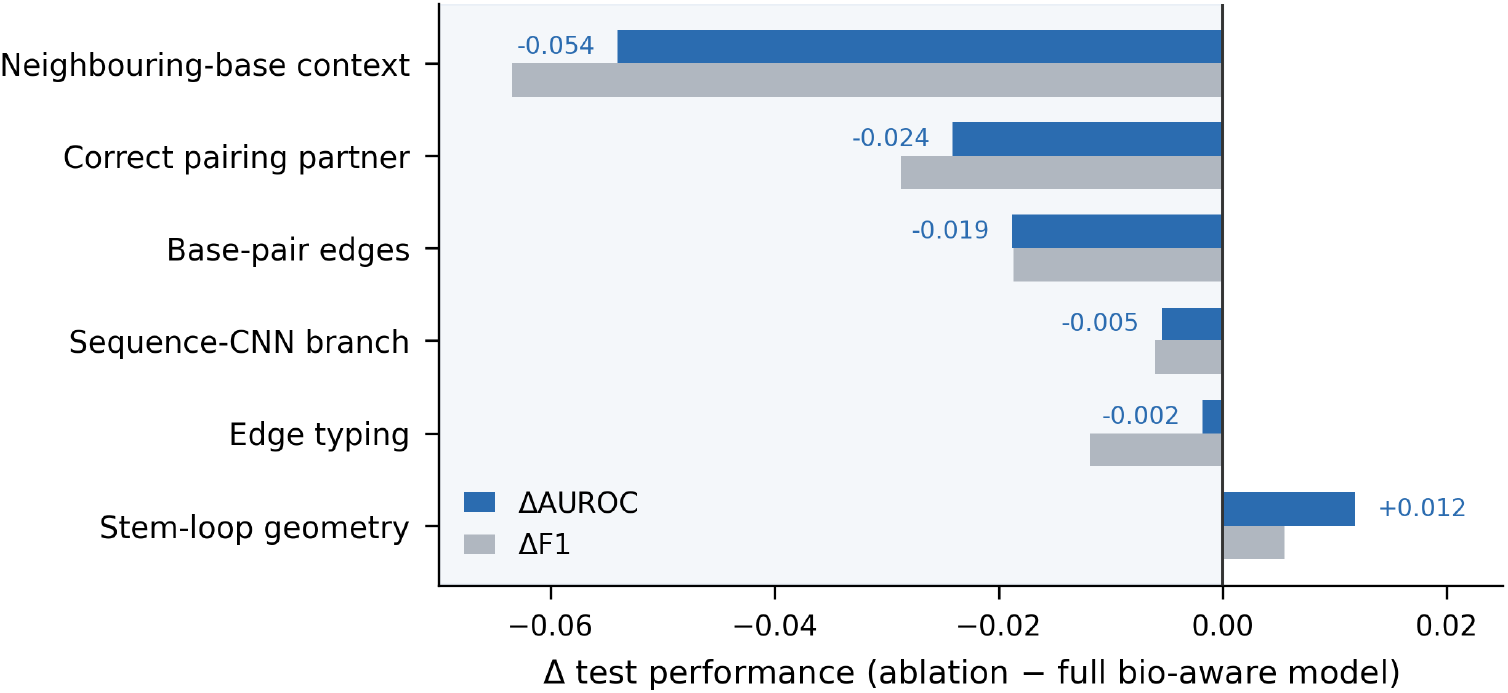
Component ablation of the bio-aware GNN on the Liver held-out test set. Change in test F1 (grey) and AUROC (blue) after each model component was individually ablated and the resulting model retrained, relative to the full bio-aware model (F1 0.854, AUROC 0.910). Bars are ordered by ΔAUROC; negative values indicate lower performance following ablation. Ablations were: neighbouring-base context (node neighbour features removed); correct pairing partner (base-pairing edges rewired to random partners); base-pair edges (base-pairing edges removed); the sequence-CNN branch (parallel sequence branch removed); edge typing (Watson–Crick/wobble/sequence edge types collapsed to a single type); and stem-loop geometry (stem/loop/junction features removed).

**Effect of Combined supervision on joint-model performance**

**Figure S8:**
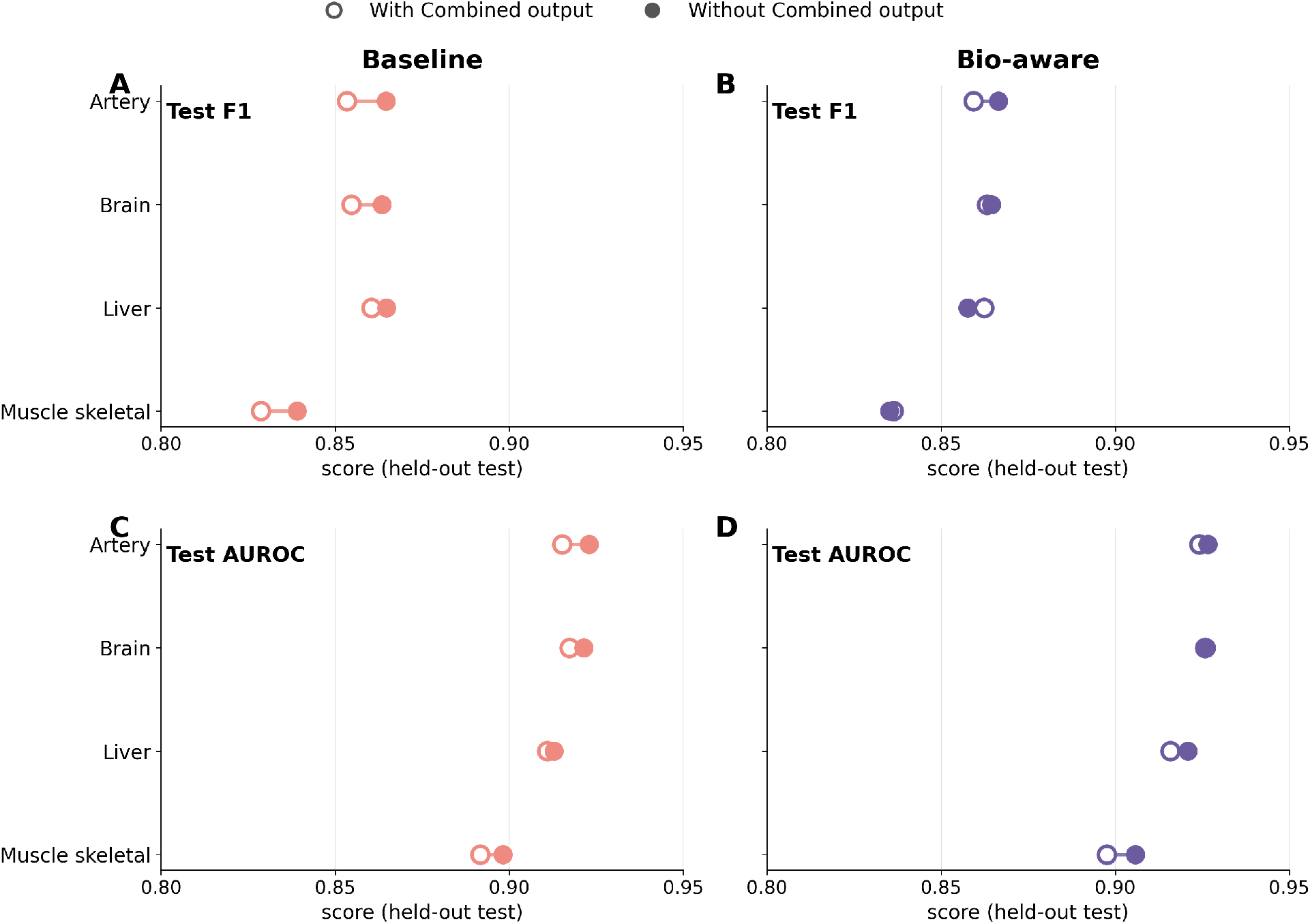
Joint-model performance with and without the Combined output. Baseline (**A,C**) and bio-aware (**B,D**) joint models were trained either with five outputs, including Combined (open circles), or with the four tissue-specific outputs only (filled circles). Models without Combined were trained independently using the same pair-disjoint split assignments and evaluation procedure. Panels show held-out test F1 (**A,B**) and AUROC (**C,D**); lines connect results for the same tissue. Across the four tissue-specific outputs, excluding Combined retained performance, with small increases in macro F1 and AUROC at the reported seed, indicating that joint-model performance on these outputs was not dependent on supervision from the aggregate Combined label.

